# Diel rhythms shape viral community structure and activity across the host domains of life

**DOI:** 10.1101/2025.10.05.680483

**Authors:** Muhammad Zain Ul Arifeen, Songze Chen, Xiaomeng Wang, Shuaishuai Xu, Jianchang Tao, Yanlin Zhao, Chuanlun Zhang, Claudia Steglich, Shengwei Hou

## Abstract

Circadian rhythms, driven by endogenous clocks and synchronized with environmental cues, are fundamental to life on Earth. While extensively studied in diverse organisms, their influence on viral ecology remains largely unexplored. This study takes the temporal dynamics of viruses in Daya Bay as an example to uncover the rhythmic control of viral replication and activity. Using a high-resolution time-series dataset collected every 2 hours over a 3-day period, we identified a total of 22,151 viral operational taxonomic units (vOTUs) and 414 Nucleocytoplasmic Large DNA Virus (NCLDV) genomes. Our analysis revealed significant diel fluctuations, with 14.48% of vOTUs exhibiting diel patterns in metagenomic abundance and 1.97% showing diel transcriptional activity. We found that these abundant diel viruses infect hosts across the domains of life, including cyanobacteria, pelagibacteria, Marine Group II (MGII) archaea and protists, all known for their diel metabolism. The expanded spectrum of host diversity with diel viral interactions significantly broadens our understanding of virus-host rhythmic dynamics in natural environmental settings. A strong positive correlation was detected between the transcriptional activities of these diel viruses and their respective hosts. Contrary to bacteriophages, which mostly peaked during the day, we demonstrated that NCLDVs showed nocturnal diel abundance with a co-fluctuating diel transcriptomic activity pattern tightly hitched to their hosts, peaking at night and declining during the day. Furthermore, we identified a rich compendium of viral genes with significant diel expression patterns, including those related to structural protein production, DNA replication, and stress response. Notably, several essential viral genes involved in stress response and repair were found to be diel transcribed for the first time, including UV-endonuclease (UvdE), peroxidase, chaperones, and early light-induced protein (ELIP). Our findings suggest that viruses across host domains actively synchronize with environmental cues to optimize their replication and transmission, despite their dependence on host metabolism. This study provides novel insights into the rhythmic control of viral communities and their intricate interactions with hosts, with profound implications for microbial community succession and biogeochemical cycles.

## Introduction

Biological rhythmicity is a fundamental mechanism that enables organisms, from bacterioplankton to mammals, to adapt to recurring environmental conditions^1,2^. Among these rhythms, the diel cycle, driven by the Earth’s 24-hour rotation, plays a particularly significant role in marine ecosystems by orchestrating biological activities that synchronize with the day-night cycle^3,4^. While diel rhythmicity has been well-studied in phytoplankton and zooplankton, which exhibit periodic changes at behavioral, physiological, and molecular levels influenced by both external factors such as light and nutrient availability, as well as internal circadian clocks^5–8^, the extent to which this rhythmicity shapes the dynamics of marine viral communities and their intricate interactions with microbial hosts largely remains unknown.

In addition to environmental oscillations, viral predation plays a critical role in shaping the spatiotemporal dynamics of marine microbial communities^9,10^. Viruses regulate bacterial populations by executing an estimated 10²³ infections per second in the ocean, significantly impacting bacterial biomass, mortality, and metabolism^11,12–17^. Due to their host-specific infections, viruses could suppress the growth and activity of target hosts, alter community composition, and thus maintain microbial diversity^18–21^. Both laboratory experiments and field studies indicate that viral activity could be synchronized with light-dark cycles and host-specific biological processes, including replication, transcription, and metabolism^3,22–24^. For example, cyanophages (viruses infecting cyanobacteria) depend on the photosynthetic activity of their hosts during the day to produce viral progeny, but predominantly lyse them at night^3,4,24^. Some cyanophages even synchronize their lytic phase with the reproduction of their *Prochlorococcus* hosts, showing higher transcriptional activity during host DNA replication^25^. This cyclic dance underscores the ability of viruses to time-track and co-opt host metabolic cycles to ensure their own proliferation while modulating bacterial populations.

Furthermore, most phages possess horizontally transferred auxiliary metabolic genes (AMGs) that are important to enhance host photosynthetic machinery and thus redirect energy from carbon fixation to nucleotide biosynthesis^26–34^. Notably, the virus-carried *psbA* gene, encoding the Photosystem II reaction center protein D1, represents a perfect example of how viruses manipulate host biological pathways to ensure their own proliferation^30,35–37^. Recently, it has been reported that viral transcription peaks during the day, while bacterial hosts show a mixed response, suggesting a complex, temporally-mediated interaction between viruses and their hosts^23,24^. In addition, various metagenomic and viromic studies have reported the prevalence of bacteriophages targeting diverse heterotrophic bacteria and some of which even show diel patterns^23,38–43^.

Despite these insights, the diel dynamics of viruses, especially those infecting abundant heterotrophic microbes in coastal waters, such as pelagibacteria and Marine Group II (MGII) archaea, remain poorly understood. Moreover, the temporal dynamics of Nucleocytoplasmic Large DNA Virus (NCLDVs) and their diverse eukaryotic hosts are not fully investigated. NCLDVs, also known as giant viruses, are a diverse group of viruses with exceptionally large genomes and virions that infect a wide range of eukaryotic hosts, including protists, algae, and even some animals^44–46^. Recent studies have begun to explore the biogeography, activity and seasonality of NCLDVs in marine systems, suggesting their potential ecological roles for biogeochemical cycles^47–49^. However, detailed investigations into their diel dynamics and ecological effects are still emerging. Studying the diel fluctuations in viral populations, including bacteriophages and giant viruses, and their influence on host metabolism could provide new insights into their roles in shaping microbial communities and related biogeochemical processes.

To address these gaps, we conducted bi-hourly time-series sampling over three days in the subtropical Daya Bay, located in the northern South China Sea. Using metagenomic, viromic, and metatranscriptomic sequencing, we tracked the temporal abundance and transcriptional activity of predominant dsDNA viruses. By linking viral dynamics with pronounced microbial diel rhythms revealed in our previous study^50^, we examined diel variations of viral genes potentially involved in host interactions, with a focus on light-dependent processes and their potential ecological roles. This study identified circadian rhythms in diverse marine viruses, particularly cyanophages, pelagiphages, magroviruses, and NCLDVs that infect cyanobacteria, pelagibacteria, MGII archaea, and eukaryotic microbes, respectively. These diel viruses possess a time-dependent regulatory mechanism that enables them to interact with their hosts and respond to environmental conditions. The results shed light on viral diel patterns and their ecological implications, providing a comprehensive understanding of how viral dynamics influence microbial community structure and function in marine environments.

## Results and discussion

### Diel dynamics orchestrate viral community structure in a coastal marine environment

To explore the temporal structuring of viral communities in the subtropical Daya Bay, northern South China Sea, we conducted a high-resolution time-series sampling every two hours over a three-day period. Surface seawater samples were sequentially filtered to capture the cellular (0.2 µm filter) and viral (0.02 µm filter) size fractions. Shotgun metagenomic and metatranscriptomic sequencing were performed on the cellular fraction, while shotgun metagenomic sequencing (i.e., viromic sequencing) was applied to the viral fraction. Viral contigs from the metagenomic assemblies of both fractions were clustered at 97% average nucleotide identity (ANI), a threshold corresponding to species delineation^51^, yielding 22,151 viral Operational Taxonomic Units (vOTUs) (**Figure 1a**; **Supplementary Dataset 1**). Of these, 68.4% were classified into viral families, with Myoviridae being the most dominant, followed by Podoviridae and Siphoviridae. The majority of classified viruses (70% of which had a predicted host), target bacterial hosts, primarily Proteobacteria and Firmicutes. A smaller proportion of viruses were predicted to infect archaeal hosts, including Thermoplasmatota, Nanoarchaeaota, and Methanobacterota (**Figure 1b**; **Supplementary Dataset 1)**. This taxonomic distribution is consistent with the established dominance of bacteriophages in coastal marine environments^23,52,53^.

**Figure 1.**
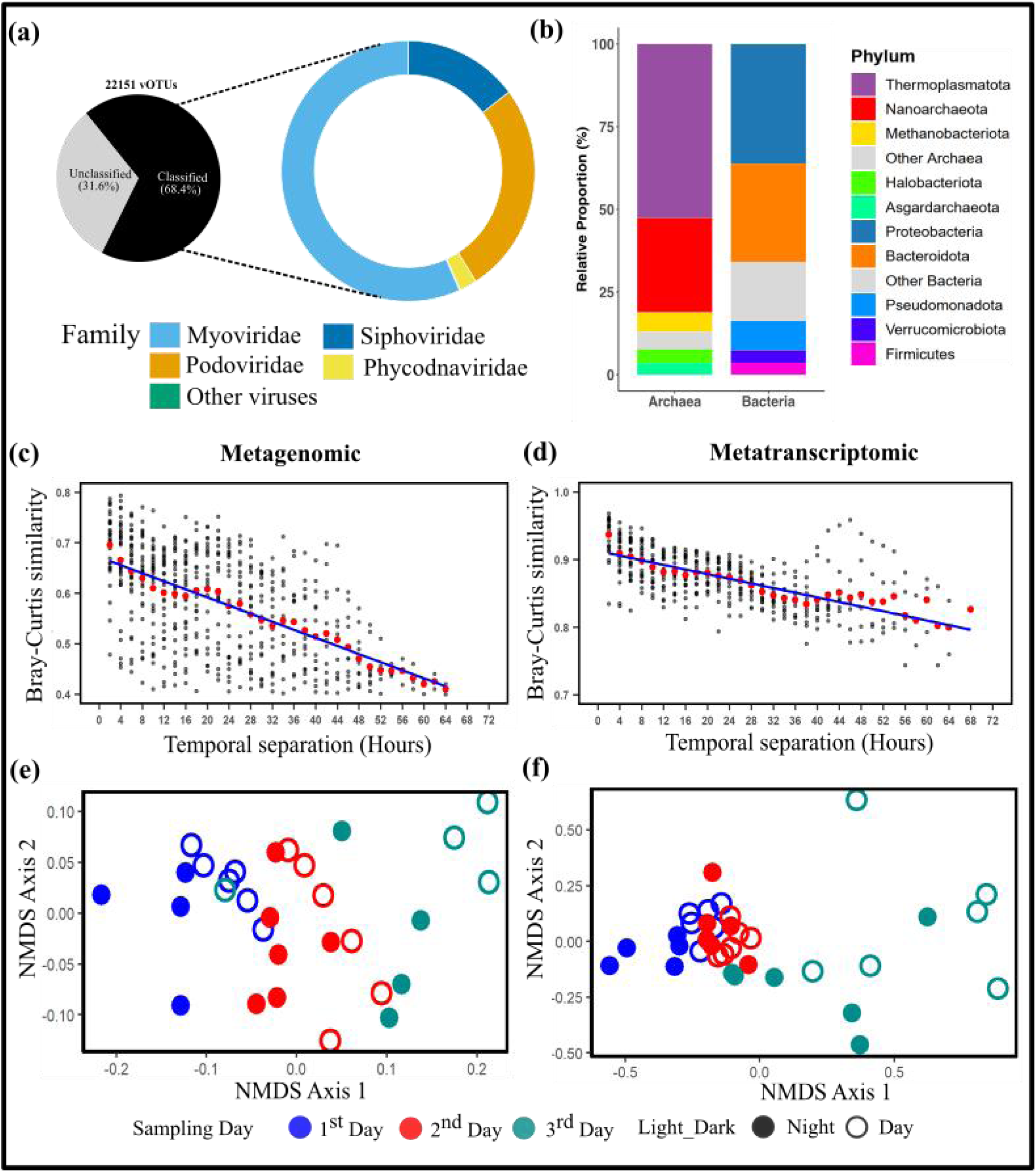
Temporal dynamics of viral communities at Daya Bay. a) The overall composition of viral community. b) Relative proportion of the predicted host for the Daya Bay viruses. (c-d) Pairwise similarity between samples based on temporal separation, shown for the metagenomic and metatranscriptomic levels, respectively. Red points indicate the mean similarity for a given temporal separation. (e-f) Non-metric Multidimensional Scaling (NMDS) ordination visually representing the temporal shifts in viral community composition.

Despite a relatively stable overall family-level composition across the study period (**Figure S1a**), temporal analysis of viral abundance revealed significant diel variations within viral communities. Peak community similarity (Bray-Curtis) occurred at 24-hour intervals, and non-metric multidimensional scaling (NMDS) analysis demonstrated distinct clustering patterns correlated with sampling days and light conditions (**Figure 1c-f**). These findings highlight the strong influence of diel cycles on viral community assembly, regardless of daily shifts in community composition. Notably, the dominant Myoviridae family, especially those infecting cyanobacteria, exhibited a pronounced diel pattern with peak relative abundance during daylight hours, consistent with the well-documented light-dependent replication of cyanophages linked to host photosynthetic activity^23,54^. In contrast, Podoviridae and Siphoviridae maintained more stable abundances, suggesting alternative infection strategies or host interactions less directly coupled to the light cycle.

We also identified representative genomes of NCLDVs infecting microbial eukaryotes. These consisted primarily of Phycodnaviridae (414 bins), with taxonomic breakdown including Algavirales (45%), Imitervirales (31.8%), Pimascovirales (8.9%), Asfuvirales (7.7%), Pandoravirales (6%) and Chitovirales (0.24%) (**Figure S1b; Supplementary Dataset 2**). Similar to bacteriophages in this study, the NCLDV community showed significant diel patterns, with Bray-Curtis similarity peaking every 24 hours and NMDS revealing distinct clusters driven by sampling day and light conditions (**Figure S2**). Interestingly, the NCLDVs displayed an inverse diel pattern with higher relative abundance during the night. This may be related to the daily vertical migration of eukaryotic phytoplankton and protists or may suggest that these viruses benefit from nighttime replication, potentially minimizing UV damage and optimizing replication within a more stable host environment^48,55^. The observed diel patterns in NCLDVs are particularly interesting given the emerging understanding of their ecological roles in marine ecosystems^45,47^. These findings underscore the profound influence of diel rhythms on viral community dynamics across host domains, revealing that different viral populations occupy distinct temporal niches in coastal marine environments.

### Phylogenetic proximity determines viral temporal dynamics

Nonparametric rhythmicity analysis using RAIN (see Methods) identified that 14.48% (3,209 vOTUs) of the viral populations in Daya Bay exhibited significant diel fluctuations (corrected P < 0.05; **Supplementary Dataset 1**). To ensure robust and reliable downstream analysis, we focused on a refined dataset of 233 diel vOTUs (1.05%) **(Supplementary Dataset 1),** by applying a more stringent significance threshold (corrected P < 0.01; see Methods) and excluding those with significant circatidal periodicity that synchronized with the tides in 12-hour cycles. Most (55%) of these viruses showed a clear peak in abundance during the day throughout the sampling duration (**Figure S3a**). The majority (61%) of diel vOTUs belonged to Myoviridae, followed by Podoviridae and Siphoviridae, consistent with the established predominance of rhythmicity in Myoviridae within marine ecosystems^23^. Notably, 45 vOTUs were identified as pelagiphages infecting *Pelagibacter* (SAR11 clade), the most dominant bacteria in Daya Bay^50^, while 5 vOTUs were cyanophages targeting *Synechococcus* or *Prochlorococcus*. Surprisingly, 20 diel vOTUs were found to infect archaea, including Thermoplasmatota and Nanoarchaeota, highlighting the broad host spectrum and ecological processes that viral circadian rhythms could impact in marine ecosystems.

Clustering of the diel abundance profiles at the metagenomic level revealed four major super-clusters (**Figure 2**; **Supplementary Dataset 1**). Clusters 1 and 2, comprising 129 vOTUs, peaked during the day (8:20-16:30), and were predominantly composed of Myoviridae targeting light-utilizing microorganisms, such as *Prochlorococcus*, *Synechococcus*, *Parasynechococcus*, *Pelagibacter*, and MGII archaea. This synchronization suggests viral replication is closely tied to host metabolic activities^23,52^. Conversely, clusters 3 and 4, comprising 104 vOTUs, peaked at night (20:30-5:20) and were dominated by Podoviridae and Siphoviridae, which potentially infect heterotrophic bacteria such as *Poseidonibacter*, UBA1924, *Pseudoalteromonas*, and SAR86. Unlike the hosts in daytime clusters, the majority of the predicted hosts in nighttime clusters lacked significant diel rhythms, suggesting that viral activity may be driven by other environmental factors, such as organic substrate or nutrient availability^50^.

**Figure 2.**
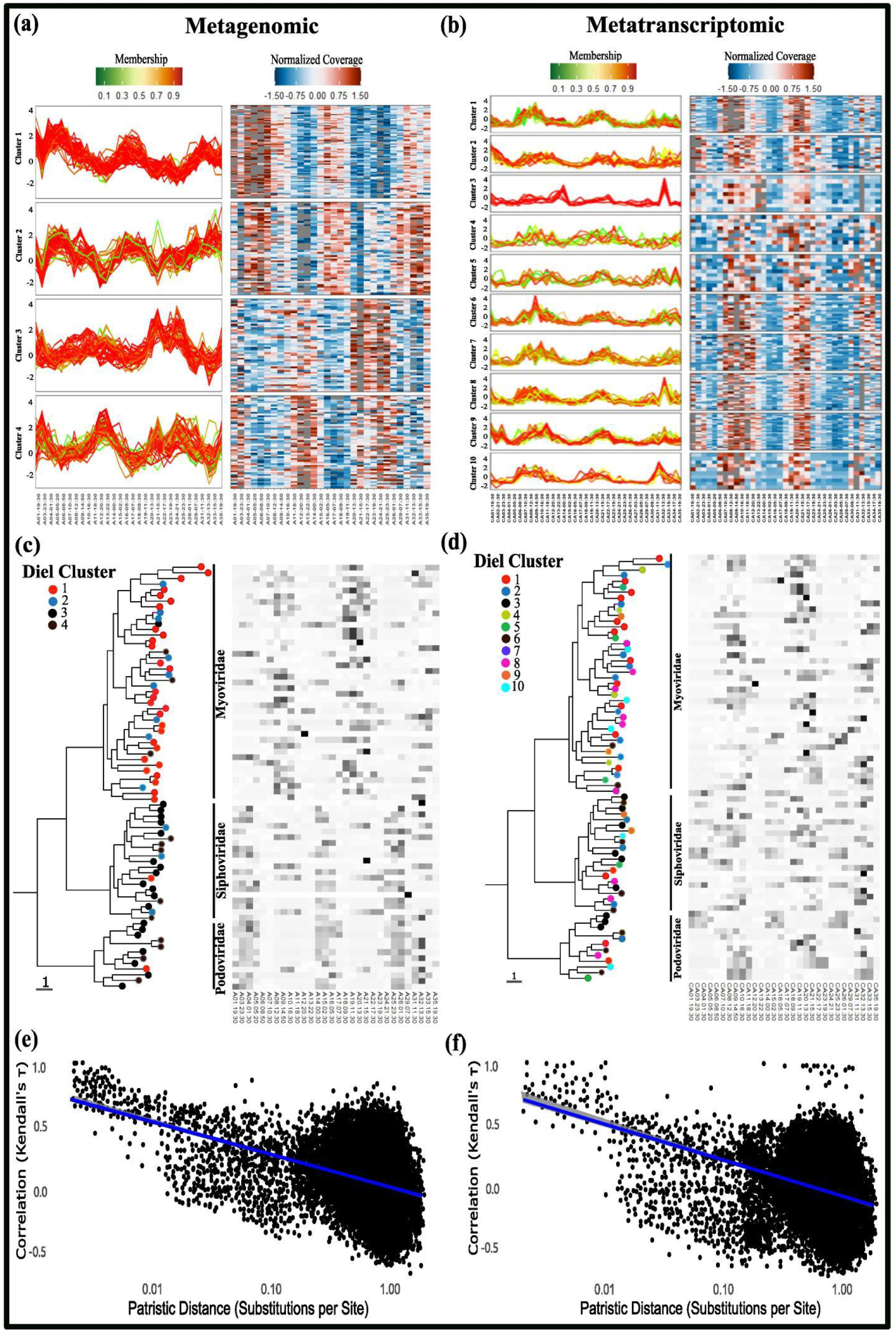
Closely related viral groups exhibit similar diel patterns. a-b) Clustering analysis of the vOTU abundance profiles using the fuzzy c-means algorithm (Mfuzz) defined four super-clusters (1–4) at the metagenomic level (a) and ten super-clusters (1–10) at the metatranscriptomic level (b). The x-axis represents sampling time, and the y-axis represents log_2_-transformed, normalized abundance. c-d) Phylogenetic analysis of the viral *g23* gene is shown alongside the mean diel abundance profiles for each gene at metagenomic (c) and metatranscriptomic (d) level. e-f) Pairwise correlations in viral abundances across time are plotted against patristic distances computed from the phylogeny in (c) and (d).

At the metatranscriptomic level, 437 vOTUs (1.97%) exhibited significant diel fluctuations (corrected P < 0.05; **Supplementary Dataset 1**). Applying a similar stringent filter (corrected P < 0.01) yielded a set of 204 vOTUs (0.92%) for further analysis, with the majority showing peak activity during the day (**Figure S3b**). Among these, 21 vOTUs were pelagiphages infecting *Pelagibacter*, and 5 were cyanophages infecting *Synechococcus* or *Prochlorococcus*. Notably, 15 vOTUs were predicted to infect MGII, suggesting diel viral activity extends beyond bacteriophages to viruses infecting archaea. These viruses are close relatives of magroviruses (within the order Magrovirales; **Figure S4c**), a group of archaeal viruses identified through metagenomic analysis with a proposed host of MGII^56^. Magroviruses are widespread in both open ocean, coastal and brackish waters, ranking as the third most abundant marine planktonic viruses after cyanophages and pelagiphages in some regions^56,57^. Similar to metagenomic abundance profiles, 10 distinct super-clusters could be revealed based on metatranscriptomic abundance profiles of vOTUs, with most of these clusters peaking between 12:30 and 16:30, but showing unique temporal dynamics. Cluster 3, which peaked later (17:30–00:30), was dominated by Siphoviridae and Podoviridae, mostly infecting alphaproteobacteria (**Supplementary Dataset 1**). A comparative analysis of metagenomic and metatranscriptomic profiles revealed 14 shared diel vOTUs, primarily targeting *Pelagibacter*, *Synechococcus*, and MGII archaea. This emphasizes that these viruses were not only present but also transcriptionally active, with their activity finely tuned to environmental and host cues.

No NCLDVs were found to exhibit diel patterns at the metatranscriptomic level using our stringent significance threshold. Therefore, we relaxed this threshold from a corrected P < 0.01 to P < 0.05 for these viruses, identifying 110 NCLDV bins with significant diel patterns (corrected P < 0.05; **Supplementary Dataset 2**) at the metagenomic level, and 16 at the metatranscriptomic level. While this relaxed threshold necessitates cautious interpretation, these observations reveal potential diel activity within this group. The 110 metagenomically abundant NCLDV bins clustered into five distinct super-clusters, with Clusters 1, 3, and 5 peaking during the night, and Clusters 2 and 4 peaking during the day (**Figure 3a; Supplementary Dataset 2**). At the metatranscriptomic level, the 16 diel NCLDV bins formed three super-clusters, where Clusters 1 and 3 peaked during the day, and Cluster 2 peaked during the night (**Figure 3b; Supplementary Dataset 2**). A complete phylogenetic tree of NCLDVs observed in this study is presented in **Figure 3c**.

**Figure 3.**
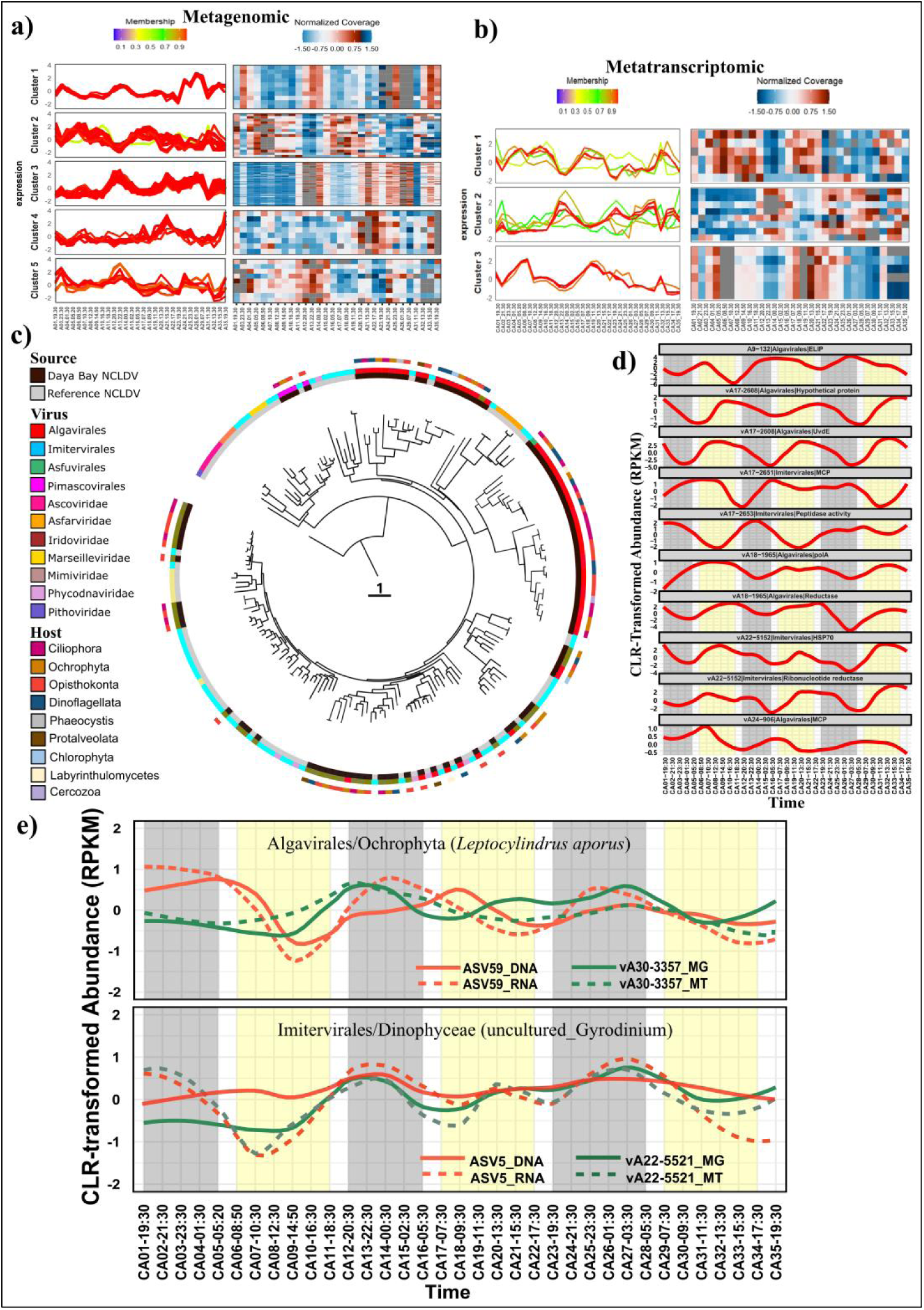
Daya Bay NCLDVs diel dynamics. Clustering analysis predicted five (clusters 1–5) and three (clusters 1-3) super-clusters defined by Mfuzz cluster analysis at the metagenomic (a) and metatranscriptomic (b) levels, respectively. Distinct clusters were calculated by applying the fuzzy c-means algorithm to the cluster vOTU abundance profiles. The x-axis represents sampling time, while the y-axis represents log_2_-transformed, normalized abundance. (c) Phylogenetic tree of NCLDVs (n = 100 from this study, plus 88 similar NCLDVs from different marine sources) based on major capsid protein (MCP) sequences. (d) Temporal profiles of genes showing significant (P-value < 0.05) diel patterns. RPKM values were CLR-transformed before plotting. Gray and yellow strips represent night and day, respectively. e) Diel patterns of selected NCLDVs and their putative hosts. Top: Algavirales (vA30-3357) and its Ochrophyts host (ASV59). Bottom: Imitervirales (vA22-5521) and its Dinophyceae host (ASV5). Dashed lines represent NCLDVs and hosts abundance at the metatranscriptomic and rRNA levels; respectively, while solid lines represent viral and host diel abundance at the metagenomic and rDNA levels; respectively.

Consistent with a recent study demonstrating that phylogenetic proximity shapes viral seasonality^49^, we found a significant correlation between viral evolutionary relatedness and diel dynamics. Closely related viral phylotypes exhibited similar diel abundance and activity patterns, as shown by phylogenetic analysis of the viral *g23* gene for phages (**Figure 2c-d**) and the *polB* gene for NCLDVs (**Figure S5a-b**). Quantitatively, a significant negative correlation was observed between phylogenetic distance and the similarity of both diel abundance (phages: **Figure 2e; NCLDVs: Figure S5c)** and transcriptional activity profiles (**phages: Figure 2f; NCLDVs: Figure S5d**). This demonstrates that evolutionary relatedness is a key factor shaping the temporal niches of viral populations. Vice versa, the phylogenetic signal in diel dynamics suggests that viral temporal regulation mechanisms are evolutionarily conserved within closely related lineages, potentially reflecting shared life strategies for optimizing host exploitation and environmental adaptation. For instance, at the family level, Myoviridae predominantly exhibited daytime activity, while Podoviridae and Siphoviridae were more active at night. A similar pattern was observed within the NCLDVs, with Imitervirales showing a nocturnal dominance and Algavirales being more active during the day (**Figure S5**). This phylogenetic structuring of diel patterns suggests that temporal regulation is not simply a plastic response to environmental conditions, but rather represents a heritable trait that has been shaped by viral evolutionary history and host interactions. This intricate interplay between viral phylogeny and diel dynamics may reflect the co-evolution of viruses with their hosts, where both partners have evolved synchronized temporal strategies to optimize their fitness in dynamic marine environments. Complete phylogenetic trees for cyanophages, pelagiphages, and magroviruses observed in this study are presented in **Figure S4**.

### Diel transcriptional dynamics reveal tight virus-host coupling and distinct viral strategies

The activities of diel vOTUs and their corresponding hosts were quantified using metatranscriptomics. Metagenome-assembled genomes (MAGs) of the putative hosts were derived from our concurrently sampled and previously published study on prokaryotic diel patterns in Daya Bay^50^ **(Supplementary Dataset 1)**. Consistent with the close coupling of viral and host abundance^9,23,52^, our data revealed a general mirroring of viral transcription with the diel activity of their putative hosts (**Figure 4**). This synchronization underscores the tight interconnectedness of viral and host dynamics in marine ecosystems^58^, which is likely constrained by host metabolic and cellular processes^23,52^. Cyanophages demonstrated a particularly strong synchronization with their hosts (**Figure S6a**). For instance, the transcriptional peak of cyanophage “A29_408_Myoviridae” coincided with the peak transcription of its predicted host “Synechococcus_C sp018623675” around noon (11:30-01:30) (**Figure 4a**). Notably, the transcriptional abundance of the virus was lower than that of the host, supporting the viral seed-bank hypothesis, which posits that only a fraction of viruses are actively replicating at any given time, depending on local host availability^59^. The aggregated transcription of all diel cyanophages peaked between 10:30–12:30 (**Figure 4b**), reinforcing the active yet synchronized interplay between viruses and their hosts. In contrast, the aggregated transcription of all non-diel cyanophages showed a steady increase at night and a decrease during the day, except on day two when chlorophyll *a* concentration surged while dissolved oxygen (DO), temperature and salinity decreased, suggesting a potential water mixing event^50^ (**Figure 4b**).

**Figure 4.**
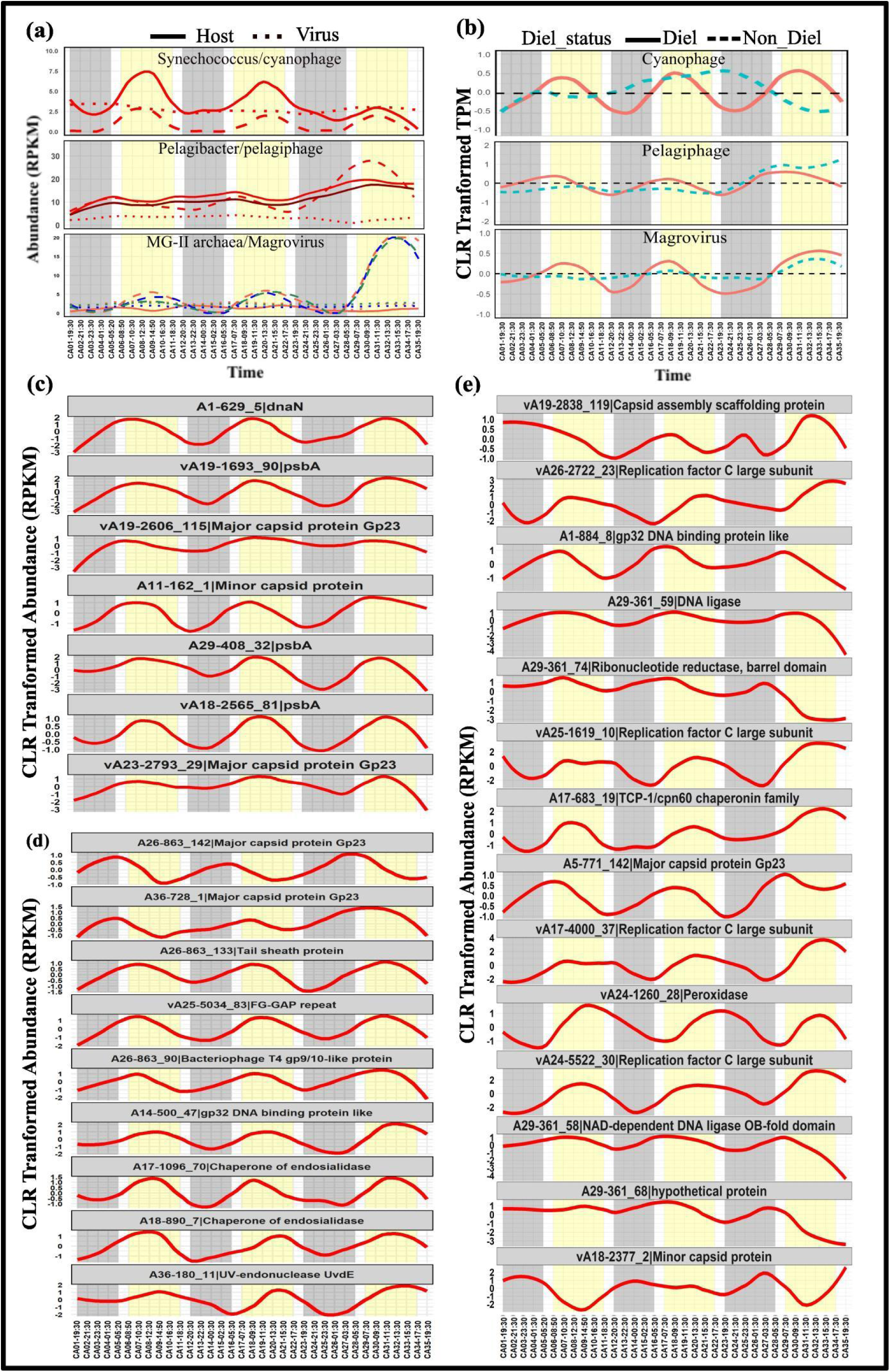
Diel synchrony of phages and their hosts activities. (a) Diel patterns of selected viruses and their hosts. Top: cyanophage (A29_408-Myoviridae) and its *Synechococcus* host (DayaUAV_MAG276/Synechococcus_C sp018623675). Middle: pelagiphage (A18_890-Myoviridae) and its putative SAR11 host (DayaUAV_MAG521/Pelagibacter sp011526385). Bottom: magroviruses and their host (DayaUAV_MAG615/MGIIa-K1). Dotted lines represent viral abundance at the metagenomic level, while dashed and solid lines represent viral and host diel abundance at the transcriptomic level. (b) Temporal profiles of aggregated diel and non-diel transcripts for cyanophage (top), pelagiphages (middle), magroviruses (bottom). Transcript Per Million (TPM) values were summed and centered log-ratio transformed (CLR). (c-e) Temporal profiles of genes showing significant (P-value < 0.05) diel patterns: (c) cyanophage genes, (d) pelagiphage genes, and (e) magrovirus genes. RPKM values were also CLR-transformed before plotting. Gray and yellow strips represent night and day, respectively.

Similarly, pelagiphages also showed apparent synchronization with their *Pelagibacter* hosts, albeit less pronounced compared to cyanophages and their cyanobacteria hosts (**Figure 4a**). For example, the pelagiphage A18_890_Myoviridae and its predicted host *Pelagibacter* sp011526385 exhibited peak transcription in the early morning (05:20-09:30) and lowest levels in the afternoon (15:30-18:30). This pattern aligns with previous findings by Martínez-Hernández et al.^60^, who reported synchronized diel transcriptional activities for pelagiphages and their putative hosts across various marine environments, with lowest transcription rates coinciding with peak solar irradiance and increased activity during early morning. The relatively weaker synchronization observed in our study may be attributed to a lower infection ratio of *Pelagibacter* cells, or the prevalence of pseudolysogenic or chronic infection states of pelagiphages^61,62^. Although the mechanisms driving pelagiphages into the pseudolysogenic state remain unclear, host metabolism or environmental cues, such as nutrient availability and light intensity, could induce the lytic cycle and promote viral production and release^52,62^. This observation supports the hypothesis that, despite their high abundance, pelagiphages may exhibit lower activity levels compared to cyanophages in coastal waters^61^. The aggregated diel transcription of pelagiphages also peaked between 07:30 and 08:30 hours, while the aggregated transcription of all non-diel pelagiphages reached their maximum expression at noon and early morning on the first and second days, respectively, but maintained elevated levels throughout the third day (**Figure 4b**).

Interestingly, viruses infecting marine planktonic archaea exhibited significant diel transcriptional fluctuations (**Figure S6b**), a phenomenon not previously reported to our knowledge. For example, magroviruses targeting MGII (e.g., vA17-4000/MGIIa-K1, vA24-5522/MGIIa-K1, and vA26-2722/MGIIa-K1) displayed peak transcriptional abundance during daylight hours (12:30-15:30), while MGII hosts showed peak transcriptional abundance at night (**Figure 4a**). This inverse relationship suggests a tightly regulated, counter-phased diel coupling, where viral activity may be optimized for periods of peak host vulnerability or specific metabolic states. Notably, the dramatically increased transcription of archaeal vOTUs on the third day coincided with higher chlorophyll *a* levels and increased DO levels^50^, suggesting a potential, albeit indirect, link between magrovirus activity and broader phytoplankton-driven ecosystem dynamics. Aggregated transcription of all diel magroviruses peaked during the day hours (09:30–13:30). In contrast, the aggregated transcription of non-diel magroviruses showed relatively uniform expression on the first two days, with elevated expression observed on the third day (**Figure 4b**).

Surprisingly, we observed a strong diel synchronization between the dominant NCLDVs, Algavirales and Imitervirales, and their putative eukaryotic hosts (**Figure 3d**). For example, Algavirales virus (vA30-3357) exhibited tightly coupled diel dynamics with *Ochrophyta* (ASV59), taxonomically assigned to *Leptocylindrus aporus*. This virus–host relationship is supported not only by our co-occurrence patterns (**Figure S7**) but also by previously reported associations in metagenomic studies^45,47^. Both viral and host transcriptomic profiles peaked during nighttime and declined during the day, reflecting an apparent nocturnal synchrony. Similarly, Imitervirales virus (vA22_5521) exhibited diel transcriptional dynamics synchronized with its putative host, *Dinophyceae* (ASV5), which is taxonomically classified as uncultured *Gyrodinium*. This pairing is supported by our correlation-based inference (**Figure S7**) and previous studies involving cultured isolates and metagenomic analyses^47,63^. While the nocturnal transcriptional activity of NCLDVs has been documented in earlier work^48,64^, the synchronized diel patterns between these giant viruses and their eukaryotic hosts at the transcriptomic level represent a novel finding. The inverse diel pattern observed between NCLDVs and bacteriophages may reflect ecological differences in their respective hosts. NCLDVs infect protists and other microeukaryotes, many of which undergo daily vertical migrations, diving to deeper waters during the day to avoid UV damage and returning to the surface water at night to feed^47,65,66^. The nocturnal peak in NCLDV abundance may therefore reflect the increased availability and activity of their hosts during nighttime hours. Alternatively, the nocturnal cycling of NCLDVs may represent an adaptation to minimize UV-induced damage to their large genomes. By limiting their replication and release during nighttime, NCLDVs may reduce their exposure to UV radiation, particularly in oligotrophic waters where UV penetration is high and the potential for DNA damage is significant. These results provide new insights into the temporal coordination of NCLDV infection cycles and host activities in natural marine systems.

Overall, the above findings highlight distinct viral-host interaction strategies in Daya Bay. The observed synchronous increase in cyanophage and pelagiphage transcription with their host abundances suggests a “Piggyback-the-Winner” (PtW) strategy^67^. Conversely, the dynamics of viruses infecting MGII, where viral activity may increase in a counter-phased manner or during periods of host vulnerability, could indicate a more complex response, potentially involving a shift towards lysogeny under specific environmental conditions to persist despite host stress^68^. This suggests that diel cycles may influence the balance between lytic and lysogenic strategies across different host-virus systems. Moreover, the nocturnal synchronization of NCLDVs and their eukaryotic hosts suggests another less explored dimension of virus-host interactions. These results highlight the complexity of viral reproduction strategies and host interactions, suggesting the need for further investigation into the underlying regulatory mechanisms and ecological drivers, particularly for viruses infecting uncultured microbial hosts. Last but not least, the diel regulation of viral activities may create temporal pulses in host infection and lysis, leading to predictable fluctuations in the availability of dissolved organic matter that could contribute to the viral shunt^69,70^ and potentially influence the microbial community structure and efficiency of carbon export.

### Diel orchestration of key viral processes

Our analysis of viral gene expression patterns at the metatranscriptomic level revealed a rich tapestry of diel regulation that extends beyond metagenomic abundance fluctuations. We analyzed diel transcriptional rhythms across the viral gene pool and identified 30 dominant genes with significant diel expression patterns (corrected P < 0.03; **Figure S8**) from cyanophages, pelagiphages, and magroviruses (**Figure 4c-e**; **Supplementary Dataset 1**). These genes are involved in core viral replication processes, or are viral-encoded auxiliary metabolic genes (AMGs), highlighting the sophisticated temporal coordination of viral replication and virus-host metabolic interactions. In viruses infecting cyanobacterial hosts, genes associated with DNA replication and virion assembly consistently peaked during daylight hours, in synchrony with the host photosynthetic activity. Specifically, nucleic acid metabolism genes peaked in the morning (around 10:30 AM), followed by capsid assembly genes at noon (around 12:30 PM), broadly aligning with the expression of photosynthesis-related genes, suggesting that viral replication is tightly coupled to host metabolic rhythms, particularly to host energy production with the facilitation of viral AMGs^23,71^. Structural protein genes, including those encoding the T4-like major capsid protein Gp23, minor capsid proteins, and the capsid assembly scaffolding protein, exhibited distinct diel profiles, peaking sequentially in the early morning, at noon, and in the afternoon, respectively. This sequential expression suggests a coordinated strategy for virus assembly and maturation synchronized with host metabolic activity, potentially optimizing virion formation efficiency and structural integrity^56,57^. Similarly, DNA replication and repair genes (e.g., replication factor C large subunit, dnaN, T4-like gp23 DNA binding protein, DNA ligase) peaked between morning and noon, further supporting efficient resource utilization during host metabolic activity^23,24,56,57^. Notably, our detection of a highly abundant and widespread *dnaN* gene in a dominant Myoviridae vOTU is particularly interesting (**Figure 5**), as this gene encoding the DNA Polymerase III beta subunit (the ‘sliding clamp’), has thus far been found in only a limited number of sequenced cyanophages, such as P60^72^. The presence of *dnaN* may provide additional advantages during the synthesis of progeny viral genomes within the host cell. This strategy allows the phage to potentially override or augment the host’s own replication machinery, thereby accelerating its life cycle and maximizing virion production, especially during periods of high host metabolic activity^73,74^.

**Figure 5.**
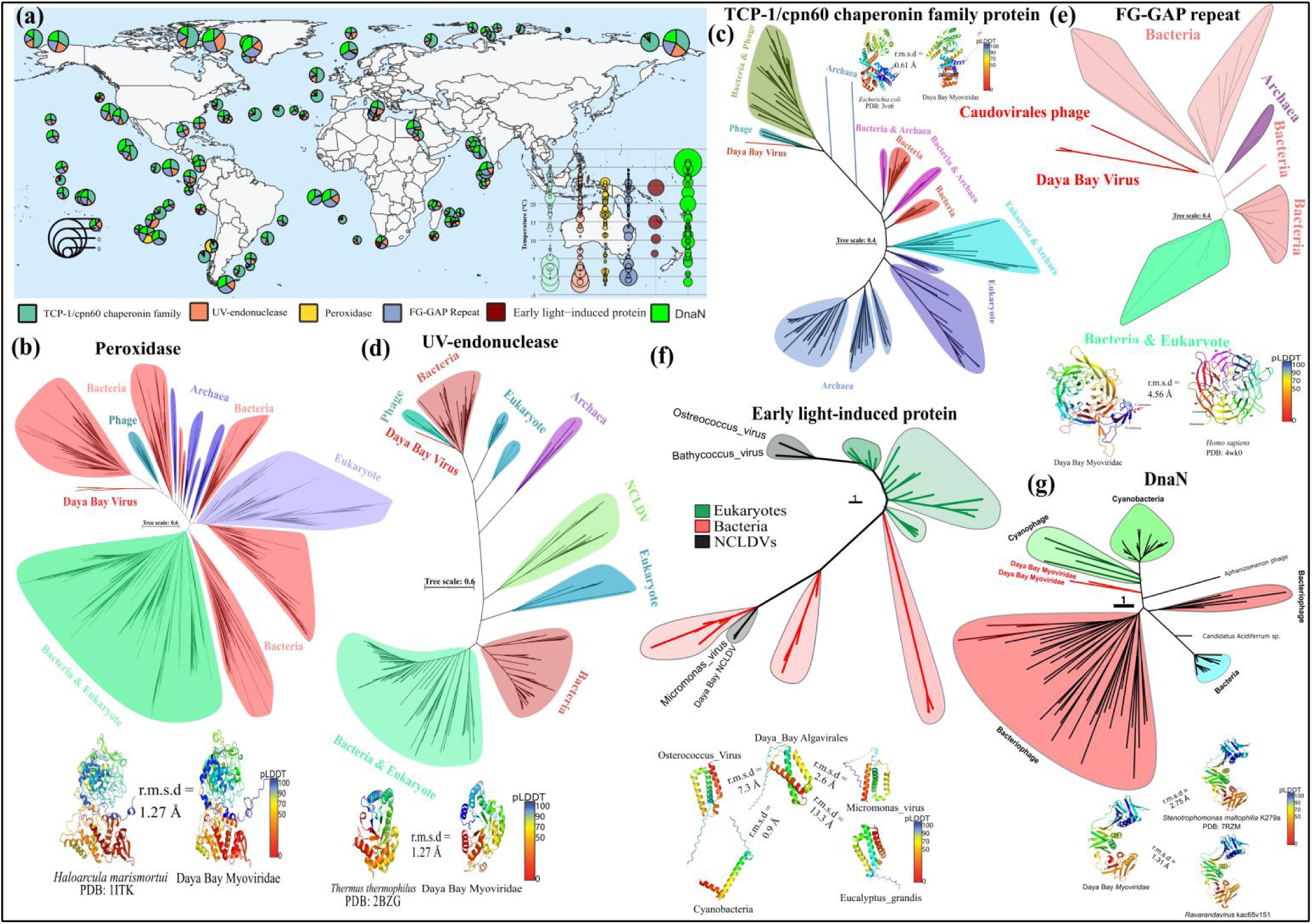
Geographical distribution, phylogenetic analysis, and structural characterization of diel-regulated viral proteins. Geographic distribution of diel genes where the normalized relative abundance of each gene is shown a). Unrooted phylogenetic trees of the peroxidase b), TCP-1/cpn60 chaperonin family c), UV-endonuclease d), and FG-GAP repeat protein e), respectively. Predicted tertiary structures of diel-regulated proteins encoded by Daya Bay viruses (Myoviridae) compared to their homologs in other organisms. Protein structures were predicted using AlphaFold3 (https://alphafoldserver.com/). Root Mean Square Deviation (RMSD) values were calculated in PyMOL to assess structural similarity between the diel proteins and their homologs. Crystal structure (left) shown alongside an AlphaFold3 predicted structure for the corresponding protein (right). These structures superpose with a root mean square deviation (r.m.s.d.) shown in the figure. The predicted structure is colour-coded by per-residue confidence scores (predicted local distance difference test (pLDDT)), as indicated in the bar.

Furthermore, genes encoding FG-GAP (phenylalanyl-glycyl, glycyl-alanyl-prolyl) repeat proteins, known for their roles in viral surface adhesion and host recognition^75–77^, exhibited peak transcription during daylight hours (**Figure 4d**). While direct evidence linking this specific gene to pelagiphage infection is lacking, its involvement in host-virus interactions via integrin receptors^78^ and its role in host surface binding in other viruses like medusavirus^79^ suggest its critical contribution to the viral targeting and infection^80^. Interestingly, one gene encoding a hypothetical protein **(Figure 4 & S9)** showed a similar diel pattern to DNA ligase genes, hinting at its potential involvement in viral genome replication, although its exact function remains to be clarified. Similarly, the ribonucleotide reductase (containing a barrel domain) illustrated unique diel transcriptional variability, with peaks shifting over the course of the sampling days (**Figure 4e**), suggesting fine-tuned regulation of nucleotide synthesis that aligns with the specific demands of viral replication and host resource availability^56^.

Additionally, we detected expression of numerous NCLDVs genes involved in diverse metabolic and cellular processes (**Figure S10a**). These included core genes crucial for viral replication (e.g., Major Capsid Protein (MCP), DNA Polymerase B (PolB) and Viral Late Transcription Factor 3 (VLTF3)), as well as various stress response genes (e.g., HSP90, HSP70, Cpn60, and UvdE). We also observed expression of genes related to cytoskeleton (actin), amino acid metabolism, lipid metabolism, and photosynthesis (Light-Harvesting Complex (Lhc)). Most of these observations were consistent with findings from previous studies on NCLDV metabolic capacities^45,48,81–83^. Intriguingly, our analysis revealed that numerous NCLDVs encode components of both glycolysis and the pentose phosphate pathway, suggesting their capacity to reprogram host central carbon metabolism. Among these expressed genes, 10 exhibited significant diel patterns (corrected P < 0.05; **Figure 3d**). These diel-cycling genes included an Early Light-Induced Protein (ELIP), UvdE, a peptidase, a reductase, HSP70, a hypothetical protein, and two core genes (MCP and PolA). Among these genes, ELIPs are particularly interesting because they are well-known for their role in protecting cellular components from damage caused by excessive light in photosynthetic organisms^84,85^ (for detail see below).

Collectively, the observed diel rhythms in viral gene expression are intricately linked to the diel cycles of host activity and environmental conditions. Viruses appear to have evolved mechanisms to synchronize their replication and gene expression with predictable fluctuations in host resource availability and environmental factors, optimizing their survival and propagation in dynamic marine ecosystems.

### Diel regulation of viral stress genes suggests phage-mediated support for host fitness

The diel regulation of viral stress genes suggests a previously unrecognized layer of phage-mediated support for host fitness in the epipelagic marine environments. Life in the sunlit zone poses significant oxidative stress due to the continuous photochemical generation of reactive oxygen species (ROS), particularly hydrogen peroxide (HOOH), from dissolved organic matter^86–88^. This threat is especially acute for microbes, including *Prochlorococcus* and *Pelagibacter*, which lack defense mechanisms needed to combat ROS, rendering them highly susceptible to HOOH-induced oxidative damage^89,90^. Consequently, these microbes depend on symbiotic “helper” heterotrophic microorganisms that are capable of detoxifying ROS from the shared environment^88,89,91^. Our findings suggest that viruses may provide a similar protective role to their hosts. We observed a distinct diel expression pattern for peroxidase genes in magroviruses, with expression peaking later in the afternoon (16:30; **Figure 4e**), likely helping their MGII hosts in scavenging HOOH. Given the continuous generation of HOOH in surface waters^86,87^, this peak expression may strategically combat accumulated oxidative damage during a period of maximal ROS production. The provision of such a defense mechanism by viruses aligns with the Black Queen Hypothesis (BQH), which posits that organisms can lose genes for essential functions if those functions are provided by other members of their community^92^. In this case, viruses may be providing a key ROS defense mechanism to their archaeal hosts. By carrying and expressing peroxidase genes, these magroviruses could be helping MGII become more of a generalist, enabling them to thrive in varied environmental conditions, such as the ROS-rich phycosphere surrounding photosynthetic organisms^93^. This phage-mediated defense provides a clear selective advantage, demonstrating a sophisticated form of viral support that shapes host ecology.

Viruses also appear to provide protection against UV radiation and general cellular stress. Our data show diel expression patterns for several stress-related genes encoding UvdE in pelagiphages and NCLDVs (**Figure 4d**), chaperones (e.g., TCP-1/cpn60 family proteins in magroviruses and chaperones in pelagiphages; **Figure 4d, e**), as well as ELIP and HSP70 in NCLDVs (**Figure 3d**), etc. The peak expression of genes encoding chaperones, heat shock proteins, and UvdE in the afternoon (14:00–15:30) coincides with peak solar radiation. This temporal regulation is consistent with their protective roles against UV-induced DNA damage and their function in facilitating protein folding and repair under conditions of temperature stress, osmotic stress and other environmental challenges^60,93,94^. The identification of ELIP genes in NCLDVs represents another novel viral-mediated stress adaptation. Its expression peaked sharply in the early morning (05:30-09:30), a period characterized by rapidly increasing light intensity. ELIPs are well-established photoprotective proteins in photosynthetic organisms, known for their role in mitigating damage from photooxidative stress caused by excessive light^84,85,95,96^. The presence of an ELIP gene in NCLDVs, and its synchronized diel expression with ambient light, suggests a sophisticated viral strategy. By helping to protect the photosynthetic machinery, viruses likely enhance the viability and sustained metabolic output of its host, which, in turn, optimizes viral replication. While viral encoding of host-like photoprotective genes is rare, giant viruses are known to encode diverse AMGs that manipulate host metabolism, including some involved in light-associated energy processes^45^. Our phylogenetic and protein structure analysis of the NCLDV ELIP gene and its homologs in eukaryotic phytoplankton further supports its viral origin (**Figure 5**), as evidenced by the distinct clustering of viral sequences into divergent clades. The global abundance distribution of ELIP-like sequences and the widespread distribution of this gene in similar NCLDVs (**Figure S10**) suggest its potential universal presence in marine giant viruses. This finding highlights a novel temporal control mechanism by NCLDVs over host photoprotection, directly impacting host cell resilience and potentially broader ecosystem productivity during critical periods of light transition. Our findings collectively demonstrate that giant viruses encode complex and diverse metabolic capabilities and may play an important role in global biogeochemical cycles through their manipulation of host metabolism.

The temporal regulation of stress genes suggests that viruses have evolved sophisticated mechanisms to respond to predictable environmental fluctuations, facilitating host survival while optimizing their own replication in dynamic marine environments. This tight synchronization and provision of environmentally relevant genes suggests a co-evolutionary strategy that optimizes the fitness of both partners, leading to the development of complex yet highly coordinated biological systems. This phage-mediated adaptation not only enhances host resilience but also influences ecosystem biogeochemistry, offering new insights into the ecological roles of viruses in marine environments.

### Global biogeography of key diel-regulated viral proteins reflects environmental selection

To further understand the environmental influences on the diel-regulated viral genes, we investigated the global biogeographical distribution of these key diel genes encoding FG-GAP repeat, peroxidase, TCP-1/cpn60 chaperonin family, DnaN, UV-endonuclease UvdE, and ELIP using the Ocean Gene Atlas (OGA) web service^97^ (**Figure 5a**). Gene encoding FG-GAP repeat protein were predominantly concentrated in the tropical Pacific and Indian Oceans with a scattered distribution in the Atlantic, suggesting adaptation to warm, oligotrophic waters. Peroxidase genes showed a broader distribution across various latitudes, notably in the Mediterranean Sea, Levantine Sea, and Indian Ocean, indicating their widespread role in ROS detoxification across diverse marine environments. The TCP-1/cpn60 chaperonin family, ELIP and DnaN genes were highly abundant in temperate, polar and subpolar regions, with the highest relative abundances in the Arctic, Antarctic, North Atlantic and Southern Oceans, highlighting their functional role in adaptation to cold, demanding environments. Conversely, UV-endonuclease genes were more abundant in tropical and subtropical regions, particularly in the Indian Ocean and the tropical Pacific, aligning with high solar UV radiation levels and emphasizing their importance in DNA repair. Previously, temperature was found to be the best predictor of the global marine plankton diversity^98^. Here, we found that the global biogeographic distribution of these diel genes was also associated with temperature (**Figure 5a**). In contrast to genes found in warmer waters, several genes exhibited a clear preference for colder marine environments. The TCP-1/cpn60 chaperonin genes and the DnaN gene both showed increased abundance at lower temperatures (below 10 °C and below 15°C, respectively), reinforcing their roles in cold adaptation and supporting viral communities in these conditions. Similarly, the ELIP gene also had its highest abundance in regions below 15 °C, suggesting its association with cold-adapted eukaryotic hosts. Conversely, the UV-endonuclease genes were more prevalent in warmer temperatures (15−25 °C), consistent with their abundance in high-UV radiation regions and emphasizing their importance in DNA damage protection. The peroxidase and FG-GAP repeat genes showed a broader distribution across temperature ranges, though the abundance of peroxidase genes increased in warmer waters, suggesting an enhanced role in ROS detoxification in such environments. These patterns suggest that viral gene repertoires are shaped by local environmental conditions, with different genes being favored in different ecological contexts, underscoring the critical role of viruses in response to marine ecosystem dynamics globally.

### Evolutionary and structural insights into diel-regulated viral proteins

To elucidate the evolutionary origins of the diel-regulated viral proteins, we performed phylogenetic analyses of these previously mentioned key genes by incorporating homologs from bacteria, archaea, eukaryotes, and other viruses. The results showed that the viral peroxidase, FG-GAP repeat, UV-endonuclease, chaperonin, DnaN, and ELIP formed distinct clades separating from their counterparts in cellular organisms **(Figure 5b-g**). This phylogenetic segregation suggests an independent evolutionary trajectory, likely driven by horizontal gene transfer (HGT) followed by adaptation to the unique constraints and host metabolism. Notably, the viral proteins exhibited closer phylogenetic affiliations to specific bacterial groups or other viral sequences, while remaining distantly related to archaeal and eukaryotic homologs. This pattern supports the idea that viruses acquired these genes through HGT, potentially from bacterial hosts or co-infecting phages, consistent with the established dynamic genetic exchange between viruses and their hosts^44,46,99^.

To gain insight into the functional implications of these evolutionary trajectories, we predicted the protein tertiary structures of these diel viral genes using AlphaFold3 (AF3) (**Figure 5**). The high confidence scores (pLDDT > 70) for the majority of residues in all predicted structures indicate the reliability of these models^100^. Comparative structural analysis with experimentally determined homologs revealed significant conservation for viral TCP-1/cpn60 chaperonin family protein (RMSD 0.61 Å), peroxidase (RMSD 0.74 Å), UV-endonuclease (RMSD 1.27 Å), DnaN (RMSD 1.31 Å) and ELIP (RMSD 0.9 Å). These low RMSD values strongly suggest that the overall protein structures are conserved and, by inference, similar functional mechanisms are expected. For the FG-GAP repeat protein, which lacks experimentally resolved prokaryotic structures, we compared the high-confidence AF3 model of the Daya Bay Myoviridae protein with the experimentally determined structure of the human homolog (PDB: 4wk0), yielding an RMSD of 4.56 Å. While higher than the other comparisons, the discernible structural similarity suggests a conserved overall folding, potentially indicative of analogous functional roles in surface interactions or host recognition. However, further experimental validation is needed to definitively ascertain its specific function in the viral context. Overall, these findings demonstrate that diel-regulated viral genes have undergone independent evolutionary adaptations while maintaining protein structural features indicative of conserved functions. This evolutionary refinement likely enables viruses to manipulate host cellular processes and optimize replication and propagation within the dynamic diel environment of marine ecosystems.

## Conclusion

Our high-resolution study of viral communities in Daya Bay shows that daily light-dark cycles have profound impacts on viral communities. These rhythms are not just abstract patterns but actively shape viral dynamics, community composition, host interactions, and gene expression across diverse viral groups infecting a broad spectrum of hosts with distinct physiology and evolutionary history. The discovery of diel cycling activities in archaea viruses and NCLDVs represents a particularly significant advance, as it reveals that even the least understood viruses are subject to temporal regulation. Notably, it was found that viral abundance and activity are often synchronized with host physiology and environmental cues, suggesting that viruses may have evolved to optimize resource exploitation, minimize environmental stress, and facilitate temporal niche partitioning within the viral assemblage. We further found that these diel regulations were extended to the transcriptional level of individual viral genes. Interestingly, genes not previously linked to circadian regulation, such as UV-endonuclease, peroxidase, DnaN, chaperones, and ELIPs, showed significant diel patterns. Altogether, these findings will significantly expand our current knowledge about how viruses respond to daily environmental fluctuations. These results argue strongly that temporal dynamics have critical importance in viral ecology. Ignoring diel rhythms risks missing a key driver of energy flow, nutrient turnover, and host-virus interactions in the ocean. Going forward, it will be important to untangle the molecular mechanisms underlying viral diel regulation and to understand how these rhythms are integrated with broader ecosystem dynamics, as well as the complex roles that viruses play in marine ecosystems in a changing ocean.

## Materials and Methods

### Sampling and environmental data collection

Time-series seawater samples were collected every 2 hours over three days (October 28-31, 2021) from Daya Bay. This high-resolution sampling was conducted using unmanned aerial vehicles (Egretta QC model EMC50 Pro, Customized, China) to autonomously collect surface seawater (∼1 m depth) from Daya Bay. During this period, sunrise and sunset occurred at approximately 5:30 and 18:30, respectively. Tidal height data was recorded, with high tides ranging from 2.0 to 2.5 meters and low tides from 1.0 to 1.1 meters^50^. Detailed information regarding sampling methodology and physicochemical parameters is available in our previous study^50^. For each time-point, 1 L of seawater samples was collected. Each sample was sequentially filtered through a 47 mm polycarbonate membrane with 0.2 μm pores (GTTP04700, Millipore, USA) and then through 47 mm aluminum oxide membranes with 0.02 μm pores (Sterlitech 1360012) using a peristaltic pump. This sequential filtration yielded two size fractions: the “cellular fraction” (> 0.2 μm) and the “viral fraction” (within 0.02-0.2 μm). All filters were immediately preserved in liquid nitrogen on site and subsequently transferred to a −80 °C freezer for long-term storage prior to DNA and RNA extraction.

### Nucleic acid extraction and cDNA synthesis

DNA and RNA were co-extracted from these filters using a modified protocol adapted from Tournier et al.^101^ and Chen et al.^102^. Briefly, filters were cut into pieces and processed using the FastDNA SPIN Kit for Soil (MP Biomedical, OH, USA) for simultaneous DNA and RNA extraction and purification. Following DNA extraction, the remaining filtrate was further processed for RNA purification. This involved DNase treatment using recombinant deoxyribonuclease I (ribonuclease-free, 2270B, TaKaRa, Shiga, Japan) to remove residual DNA, followed by purification with the RNeasy MinElute Cleanup Kit (Qiagen, Maryland, USA). Finally, complementary DNA (cDNA) was synthesized from the purified RNA using the PrimeScript™ Double Strand cDNA Synthesis Kit (TaKaRa, 6111A).

### Metagenomic, viromic and metatranscriptomic sequencing

Metagenomic, viromic, and metatranscriptomic libraries were sequenced on the DNBSEQ-T5 platform (BGI-Shenzhen and BGI-Qingdao), generating 100/150 bp paired-end reads. This yielded an average of 122.24 million reads for cellular metagenomes and 3.12 million reads for metatranscriptomes per sample. Unfortunately, the majority of the viromic sequencing reads did not meet quality standards, with only seven samples passing quality control and retained for further analysis by combining it with metagenomics data for vOTU identification. Raw reads from all sequencing libraries were trimmed and quality-filtered using fastp v0.23.1^103^.

### Viral contig identification and vOTU generation

Quality-filtered reads from both cellular and viral size fractions were independently assembled using MEGAHIT v1.2.9^104^. Contigs shorter than 1kb were removed using MetaQUAST v3 (-m 1000)^105^ and BBMap (minlength=1000) tools. The remaining contigs were combined and clustered into representative sequences at 97% identity using CD-HT v4.8.1^106^ to reduce redundancy. To identify viral sequences within the assembled contigs, a multi-step approach was employed. First, geNomad v1.11.1^107^ was used for initial viral sequence identification. Next, DeepMicroClass v1.0.3^108^ was applied to filter out any remaining non-viral sequences. Finally, seqkit2^109^ was used to retain only sequences longer than 10 kb. These refined viral sequences were then de-replicated at 97% identity with 85% overlap using CD-HT v4.8.1^106^ with parameters: -c 0.97 -aS 0.85 -d 400 -T 20 -M 20000 -n 5, generating 22,151 vOTUs. vOTU quality was assessed using CheckV v1.0.1^110^, which identified 2,122 high-quality viral genomes (completeness > 50%; contamination < 5%). Open reading frames (ORFs) within these vOTUs were predicted using Prodigal v2.6.3 with default parameters^111^.

### Taxonomic classification and host prediction of vOTUs

Taxonomic classification of vOTU was performed using a combination of approaches. First, vOTU sequences were compared to Viral Protein Family (VPF) database in IMG/VRv4.0^112^ and the ViralZone database using VPF-Class^113^ with the following parameters: --membership_ratio 0.5 --confidence_score 0.2. To further validate these annotations, Kaiju v1.9.2 l^114^ was employed to compare vOTUs against the NCBI RefSeq virus database. The lifestyles of vOTUs were predicted using VIBRANT v1.2.1^115^ with default parameters. Putative hosts for the vOTUs were predicted using iPHoP v1.3.3^116^, a software that integrates multiple methods for predicting the hosts of metagenome-derived viruses.

### Estimating vOTUs abundance

The abundance of each vOTU was estimated by mapping reads and calculating reads per kilobase million (RPKM) values. Quality-controlled reads from both metagenomic and metatranscriptomic datasets were individually mapped to the vOTUs using BWA v2.2.1^117^. RPKM values were then calculated using CoverM v0.3.1^118^ with the following parameters: --trim-min 0.10 --trim-max 0.90 -- min-read-percent-identity 0.95 --min-read-aligned-percent 0.75 -m rpkm. The resulting RPKM values, representing the relative abundance of each vOTU in each sample, are provided in **Supplementary Dataset 1**.

### Identification and analysis of viral AMGs

Viral contigs were first processed using Prodigal v2.6.3^111^ to predict genes. DRAM-v v1.2.0^119^ was then used to annotate the vOTUs and identify putative AMGs. To improve the accuracy of AMG identification, a two-step approach was employed. First, Virsorter2 v2.2.4^120^ was run, and viral contigs with a maximum score greater than 0.5 were selected. Virsorter2 was run again with the -prep-for-dramv option to generate an input file specifically for DRAM-v. In addition to DRAM-v, vOTUs were annotated using KEGG, Pfam, and VOG databases via VIBRNAT v1.2.1^115^ with default parameters to further identify potential AMGs. To minimize false positives and ensure annotation accuracy, the AMG outputs from both VIBRNAT and DRAM-v were manually reviewed^121^.

### Identification of NCLDVs

For the systematic identification of giant virus genomes within our metagenomic datasets, we employed the PIGv pipeline^122^. PIGv integrates several existing software tools and scripts to streamline giant virus discovery. As input, we used previously assembled contigs and their corresponding coverage files. The pipeline first performed metagenomic contig binning using MetaBAT2^123^. Proteins were then predicted for these bins using Prodigal-gv^111,112^, a specialized version of Prodigal trained with giant virus models and accommodating alternative genetic codes. Subsequently, bins were screened using the NCLDV markersearch script^124^ to identify key marker genes characteristic of giant virus families. Bins positively identified as NCLDVs based on these marker genes were selected for further processing. These NCLDV bins were then dereplicated at 98% average nucleotide identity using dRep v3.4.2^125^. Protein-coding regions for the dereplicated NCLDV bins were annotated using eggNOG-mapper^126^ with default settings. Taxonomic assignment for these genomes was performed utilizing GVClass^127^. To predict putative hosts for the identified NCLDVs, a co-occurrence network was constructed. This network linked NCLDVs to eukaryotic amplicon sequence variants (ASVs), which were obtained from a previously described dataset^50^, using FlashWeave^128^. The resulting correlations were then filtered using a weight cutoff of 0.4^129^ and visualized using the R package igraph v2.1.4^130^, as depicted in **Figure S7**.

### Diel periodicity analysis

To identify vOTUs and viral genes exhibiting diel periodicity, we applied the Rhythmicity Analysis Incorporating Non-Parametric Methods (RAIN) algorithm using the RAIN R package v1.26.0^131^. This analysis was performed on the relative abundance estimates of vOTUs in both metagenomic and metatranscriptomic datasets. For the metatranscriptomic data, diel periodicity was assessed at both the aggregated vOTU level and the individual viral gene level. To focus on vOTUs and transcripts with sufficient data for robust analysis, low abundance vOTUs (those with < 2 read mapping per time point, on average) were filtered out. The resulting P-values from RAIN were corrected for multiple comparisons using the Benjamini and Hochberg method^132^. vOTUs or genes with corrected P-values less than 0.05 were considered to exhibit statistically significant diel periodicity. This significance threshold aligns with established practices in diel periodicity analysis^133^. Furthermore, to account for potential circatidal influences, we employed a modified RAIN test to identify and exclude vOTUs with significant circatidal periodicity (12-hour cycles). This process identified and removed 26 vOTUs with significant circatidal rhythms (**Supplementary Dataset 1**). In addition, to account for potential biases in relative abundance measurements from metagenomic data^134,135^, we validated the rank-based results from RAIN using centered log-ratio (CLR) transformation, which is less susceptible to such biases^135^.

### Clustering analysis of diel vOTUs

To group vOTUs with similar diel patterns, we performed fuzzy clustering using the fuzzy c-means (FCM) algorithm implemented in the Mfuzz package^136^. This algorithm employs an iterative process to assign vOTUs to clusters by minimizing the objective function based on the Euclidean distance between data points (vOTU abundance profiles) and cluster centroids. The resulting cluster membership values for each vOTU indicate the strength of its association with each cluster, allowing for vOTUs to have partial membership in multiple clusters. This approach provides a more nuanced representation of vOTUs groupings compared to traditional hard clustering methods.

### Statistical analyses

All microbial statistical analyses were performed in R version 4.2.2. Alpha and beta diversity of viral communities were calculated using the vegan R package v2.6–4^137^.

## Acknowledgments

We appreciate Prof. Qinglu Zeng for his constructive comments on the manuscript. This study was supported by the National Natural Science Foundation of China (grant numbers: 42476109, 42276163, 42406144), the Shenzhen Science, Technology and Innovation Commission Program (grant number: JCYJ20220530115401003, SUSTech Education Reform Programme (grant number: SJZLGC202437), and a grant from the Deutsche Forschungsgemeinschaft (SPP 2330 project number 464976318) to C.S.

## Author contributions

S.H. and C.Z. conceived the idea and designed the experiments. S.H. and S.C. conducted field sampling. S.C. performed DNA extraction, amplicon, and metagenomic data analysis. M.Z.U.A., X.W., and S.C. analyzed the data and drew the figures. M.Z.U.A. and S.H. wrote the manuscript. S.H., Y.Z. and C.S. reviewed and revised the manuscript. All authors contributed to the final version of the manuscript.

## Ethics declarations

### Animal and human rights statement

This article does not contain any studies with human participants or animals performed by any of the authors.

### Data availability

Raw reads generated in this project have been deposited at NCBI under the umbrella project number PRJNA949377.

