## Supplementary figures for "Diel rhythms shape viral community structure and activity across the host domains of life"

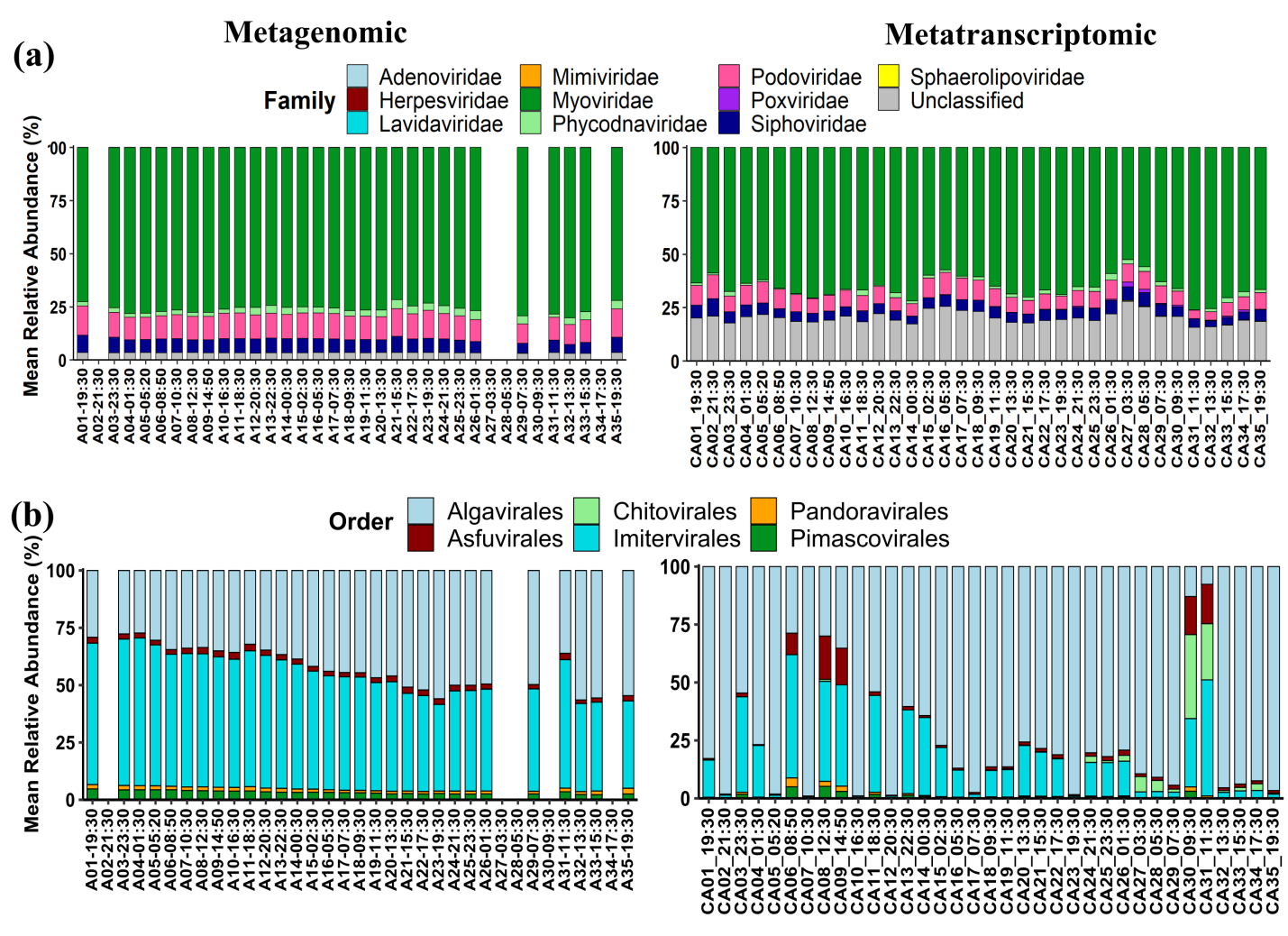

**Figure S1.** Overall viral community composition (a). Nucleocytoplasmic large DNA viruses (NCLDVs) composition (b).

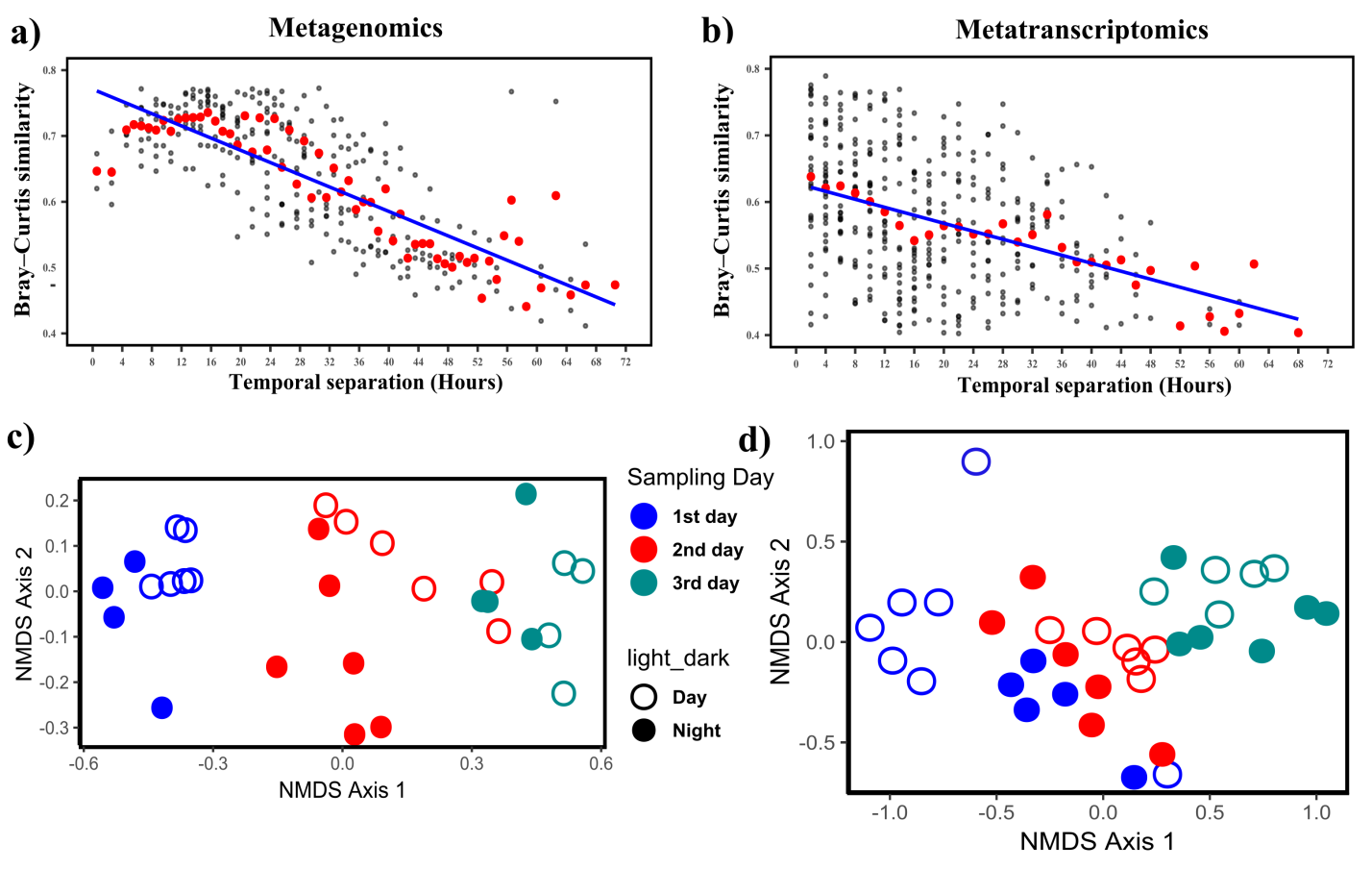

**Figure S2**. Temporal dynamics of Daya Bay’s NCLDVs. (a-b) show the pairwise similarity between samples, at metagenomics and metatranscriptomic level, respectively. (c-d) depicts the non-metric multidimensional scaling (NMDS) ordination, visually representing the temporal shifts in viral community composition, at metagenomics and metatranscriptomic level, respectively. In panels (a and b), red points indicate the mean Bray-Curtis similarity for a given temporal separation.

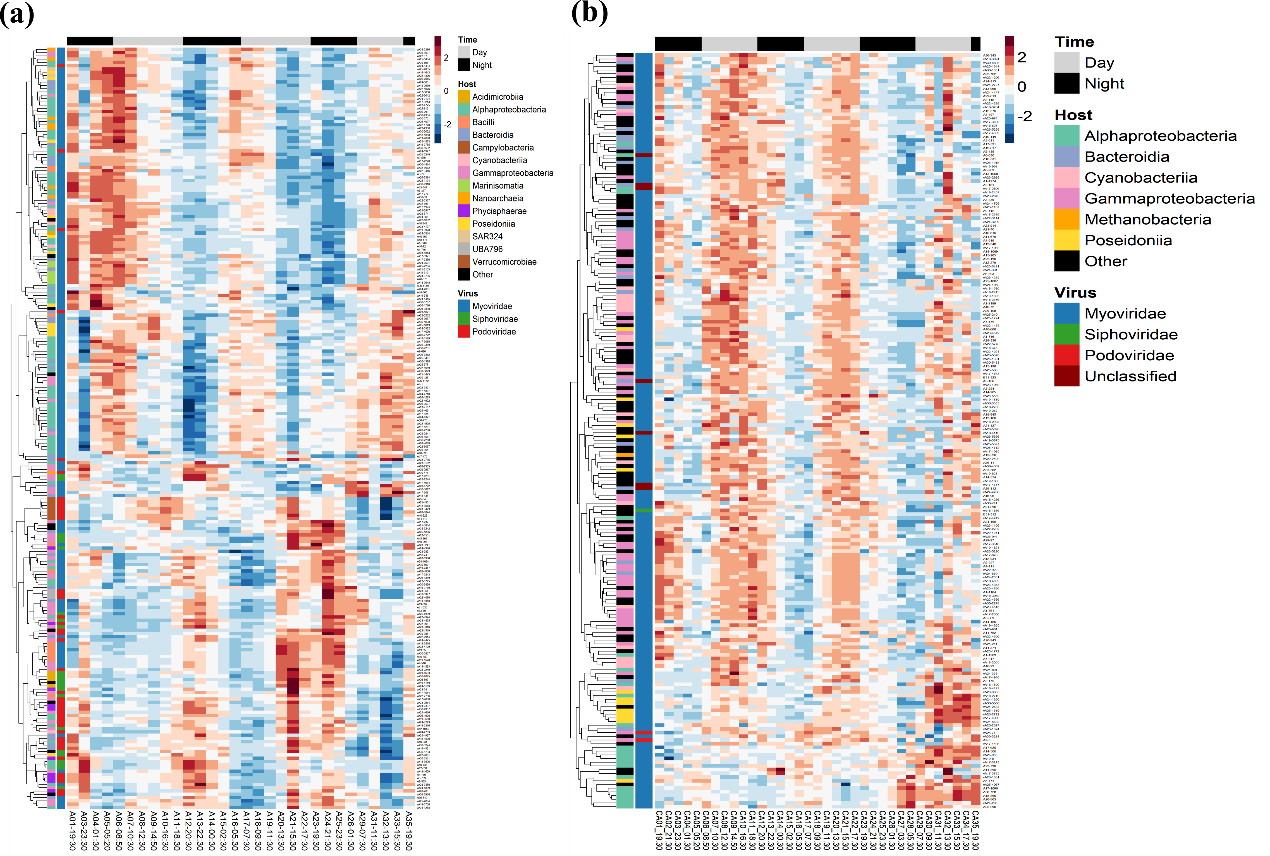

**Figure S3.** Euclidean single cluster analysis of diel periodic rhythms in vOTUs at metagenomic a) and metatranscriptomic b) level. The heatmap denotes the diel vOTUs determined by the RAIN periodic test with the corrected P-value < 0.01. All measurements were scaled to have a mean of zero and a variance of one

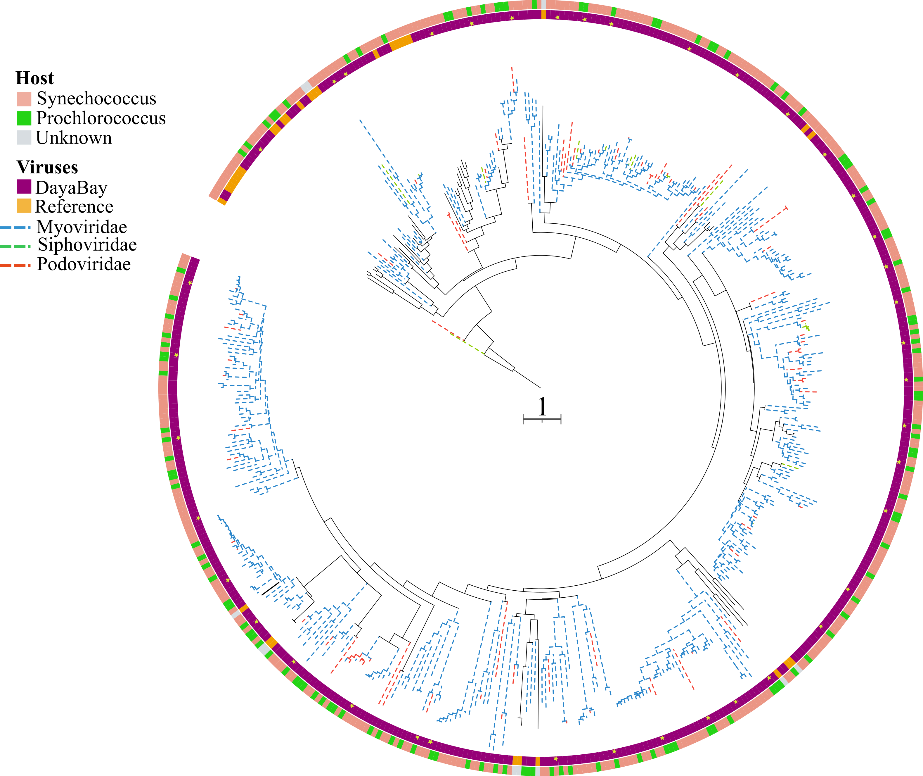

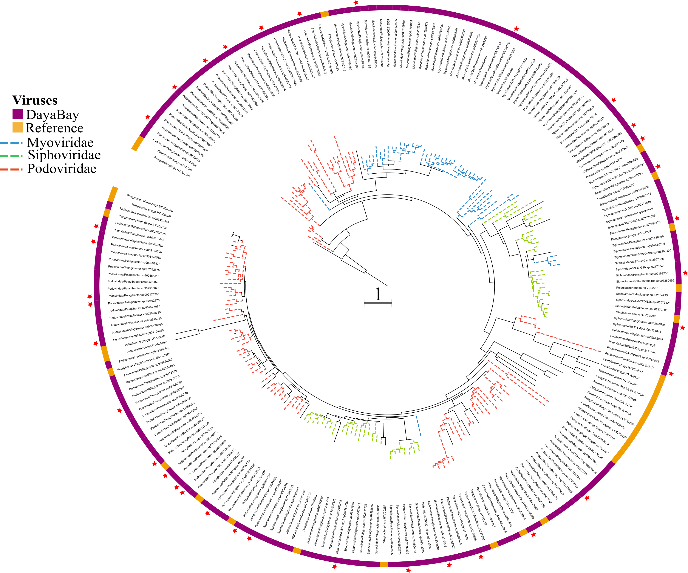

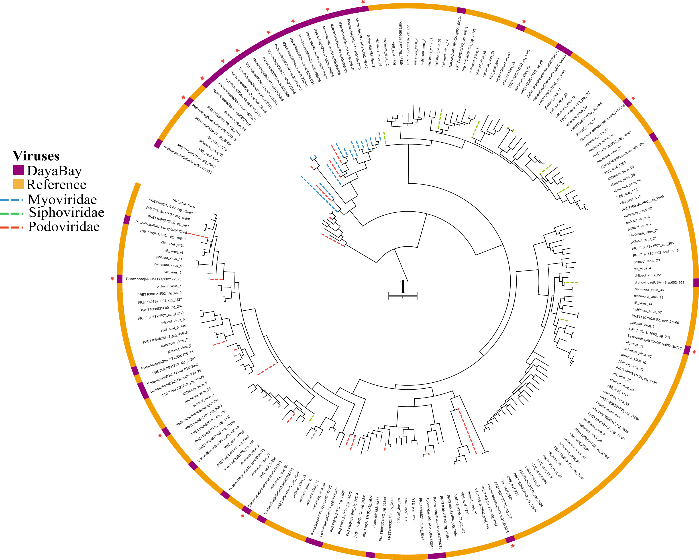

**b)**

**c)**

**a**

**Figure S4**. Unrooted phylogenetic trees for various marine phage groups were constructed as follows: (a) Cyanophages (n = 464 from this study, plus 50 similar matches from RefSeq proteins) based on large terminase (TerL) sequences. (b) Pelagiphages (n = 178 from this study, plus 35 similar matches from NCBI RefSeq sequences) based on DNA polymerase A (DNApolA) sequences. (c) Magroviruses (n = 47 from this study, plus 152 similar matches from NCBI RefSeq sequences) based on DNA polymerase B (DNApolB) sequences. Phages showing significant diel patterns were highlighted with a star sign

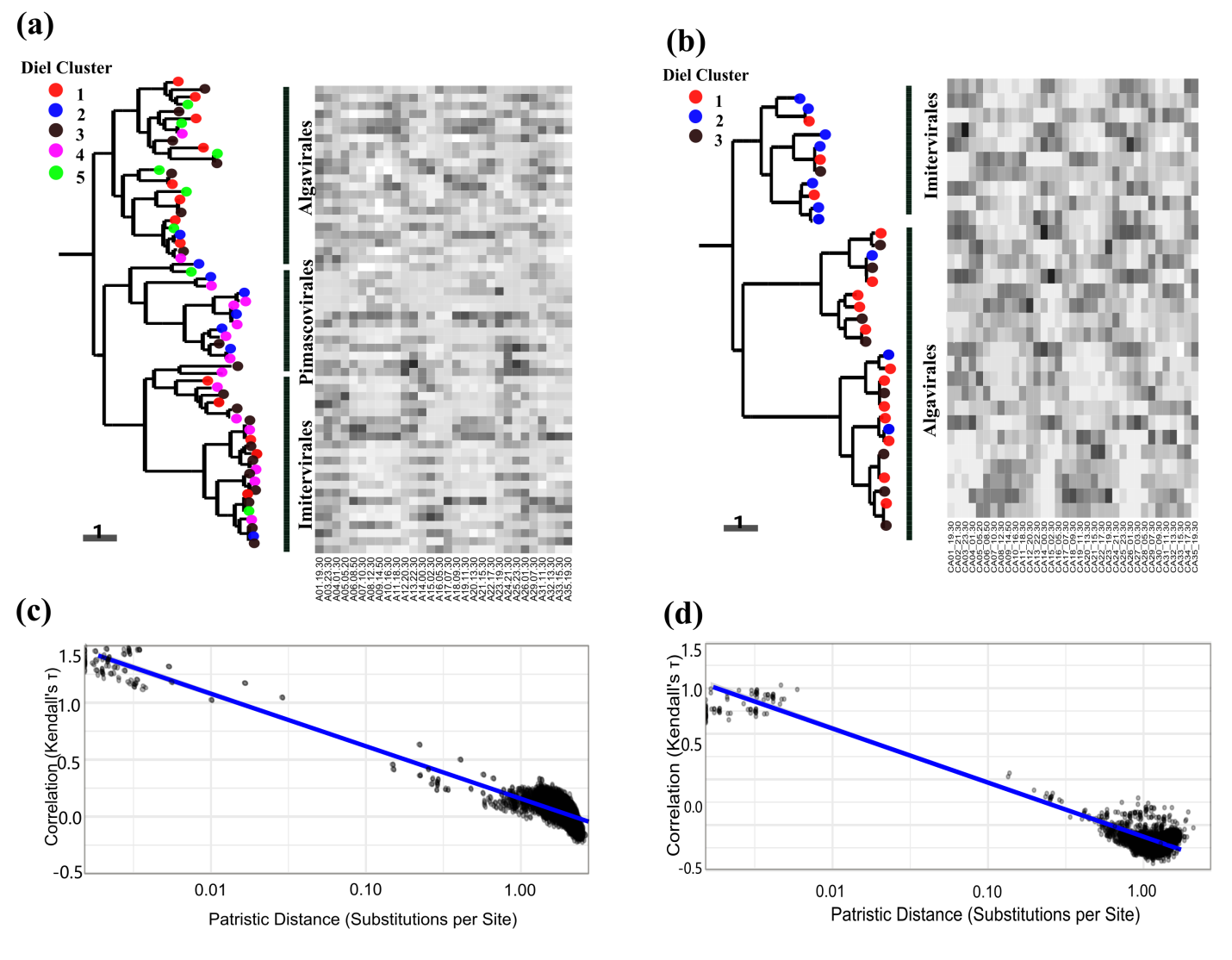

**Figure S5.** Closely related viral groups tend to have similar diel patterns. a-b) Phylogenetic analysis of NCLDVs *polb* gene and mean diel abundance profiles for the gene at metagenomic (a) and metatranscriptomic (b) level, respectively. c-d) Pairwise correlations in viral abundances across time plotted against patristic distances computed from the phylogeny in (a and b).

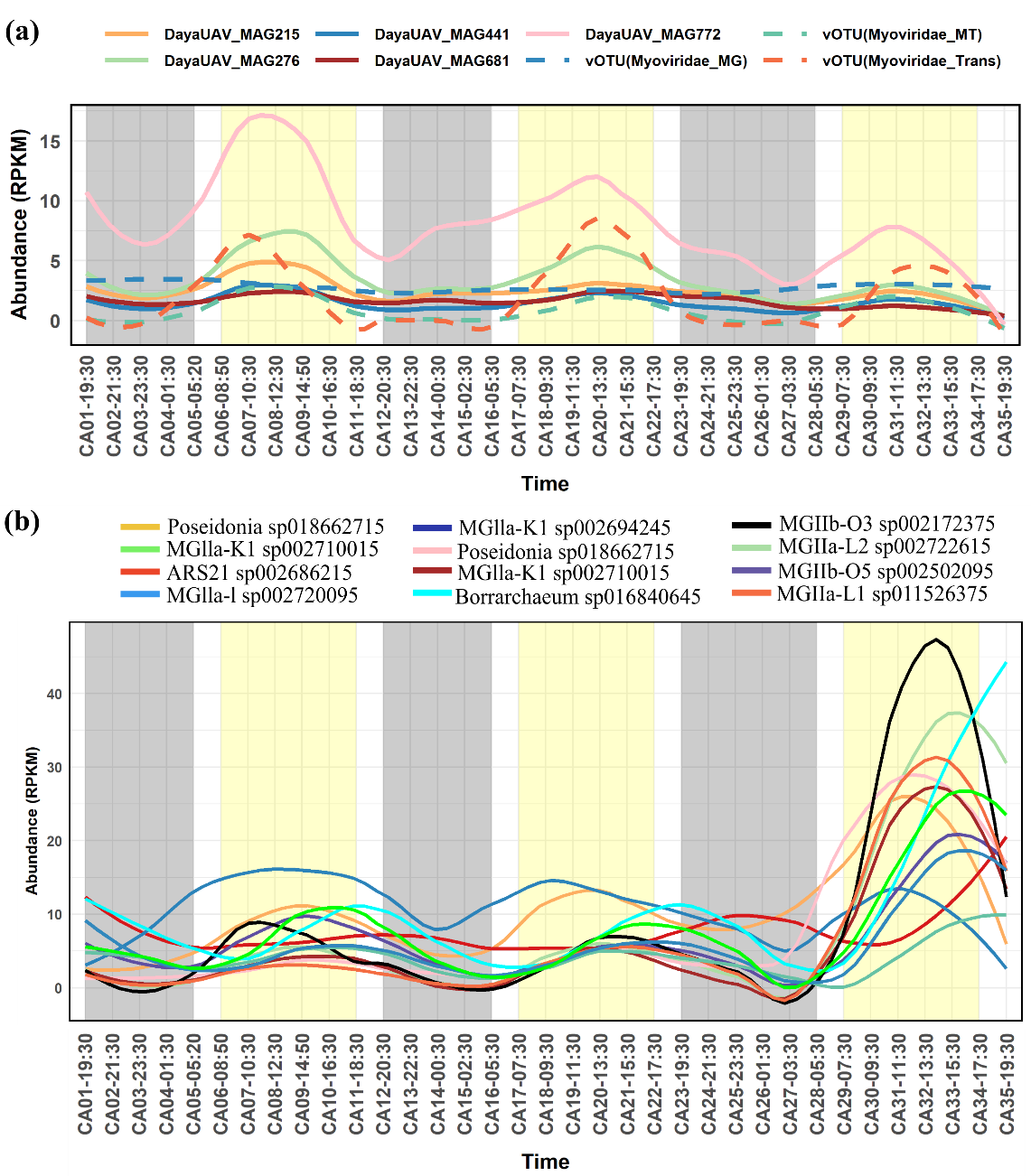

**Figure S6.** Diel transcript abundance of cyanophages and phages infecting archaea. (a) Diel transcript abundance of cyanophage and their putative hosts. (b) Diel abundance of phages infecting Thermoplasmatota. Metagenome-assembled genomes (MAGs) from our previously published study (Chen et al., 2024) and viral operational taxonomic units (vOTUs) from the current study were used. Cleaned metatranscriptomic reads were mapped to these genomic resources to determine transcript abundance.

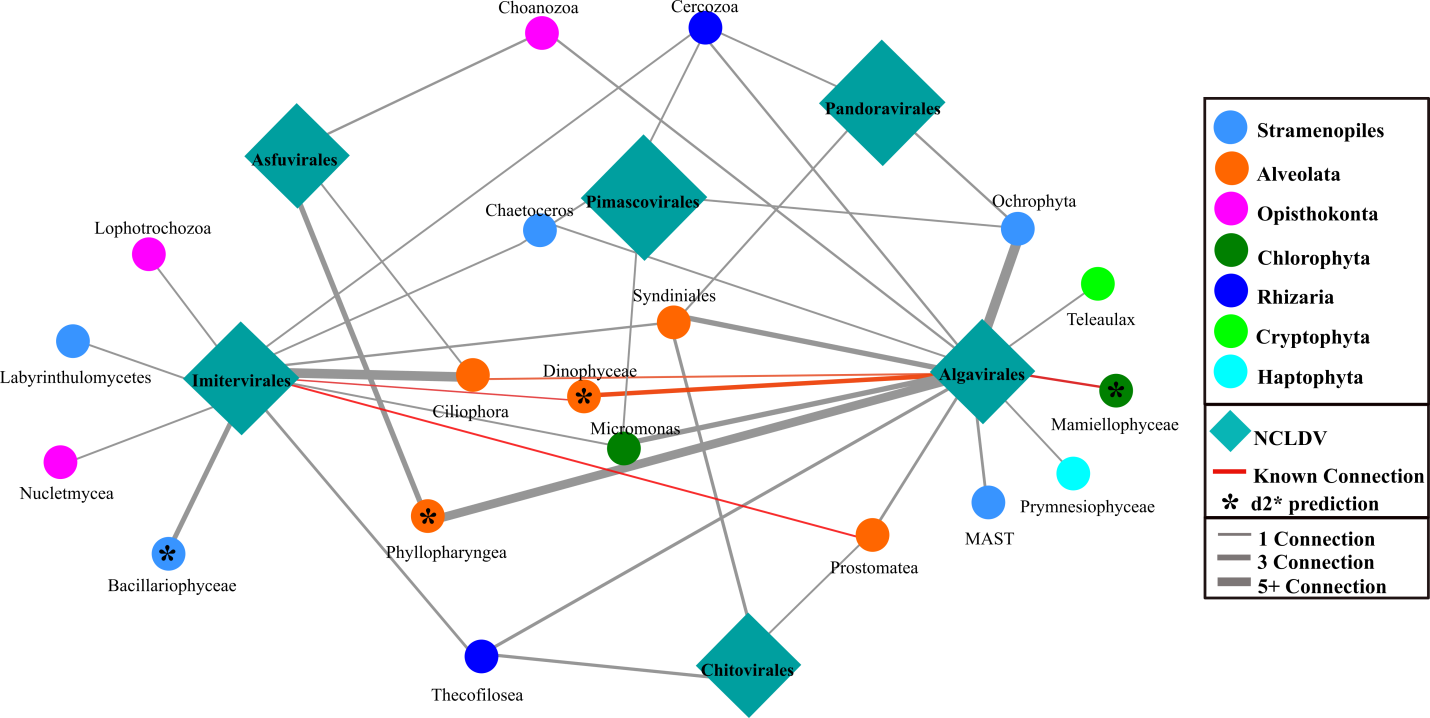

**Figure S7:** Co-occurrence network of NCLDV PolB marker gene and eukaryote ASVs using 18S data. A weight cutoff of 0.4 was used to filter connections, and the number of connections is shown by the edge thickness. Known NCLDV and eukaryotic host pairs from cultured isolates are shown in red and connections match and d2* results are shown with a star (*).

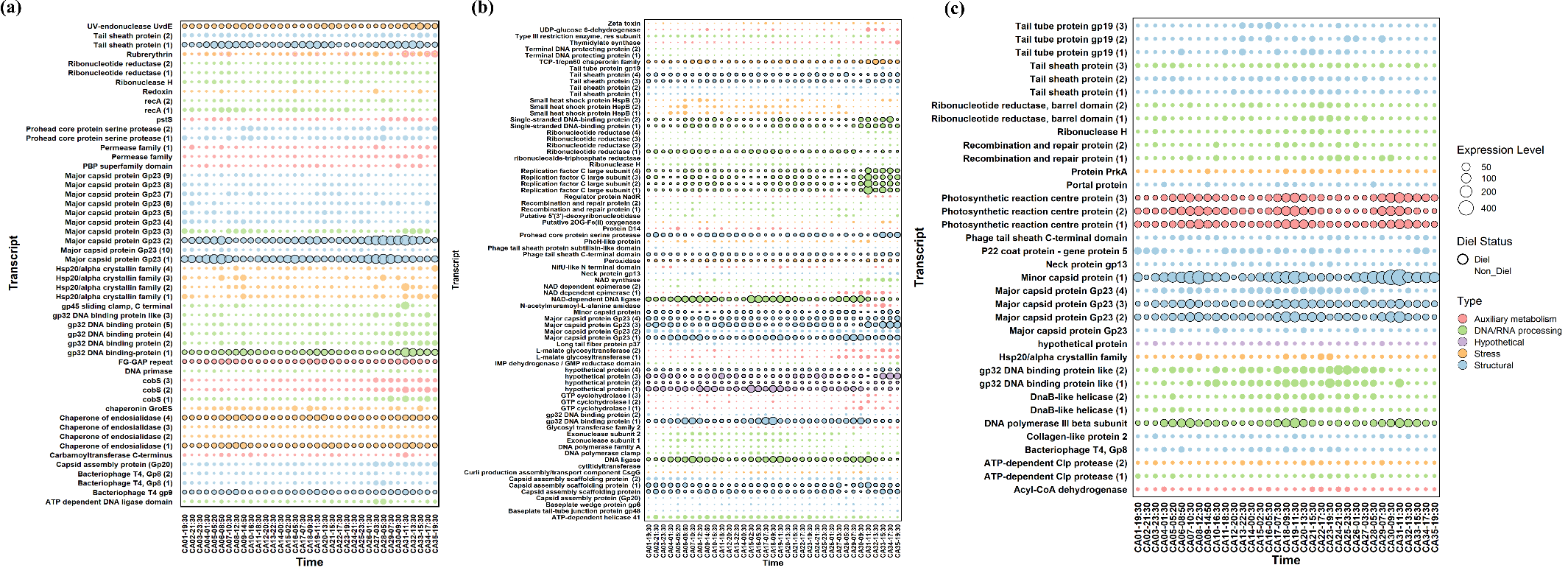

**Figure S8**. Diel expression patterns of abundant viral genes. Bubble plots illustrating the abundance of the most abundant transcripts over the diel time-course for: (a) pelagiphages, (b) magroviruses, and (c) cyanophages. Bubble size is proportional to the relative abundance of the respective viral transcripts. From all genes exhibiting significant diel patterns (corrected P-value < 0.05), we selected the top 30 most abundant genes for further analysis. All selected genes had a corrected P-value < 0.03, ensuring robust diel periodicity. Numbers in parentheses indicate the number of distinct viral contigs encoding the associated gene. While initial annotations included hypothetical proteins, further bioinformatic analysis allowed us to assign a putative function to some of these, especially one specific pelagiphage hypothetical protein (formerly designated "hypothetical protein (1)"), which is now identified as a Bacteriophage T4 gp9/10-like protein.

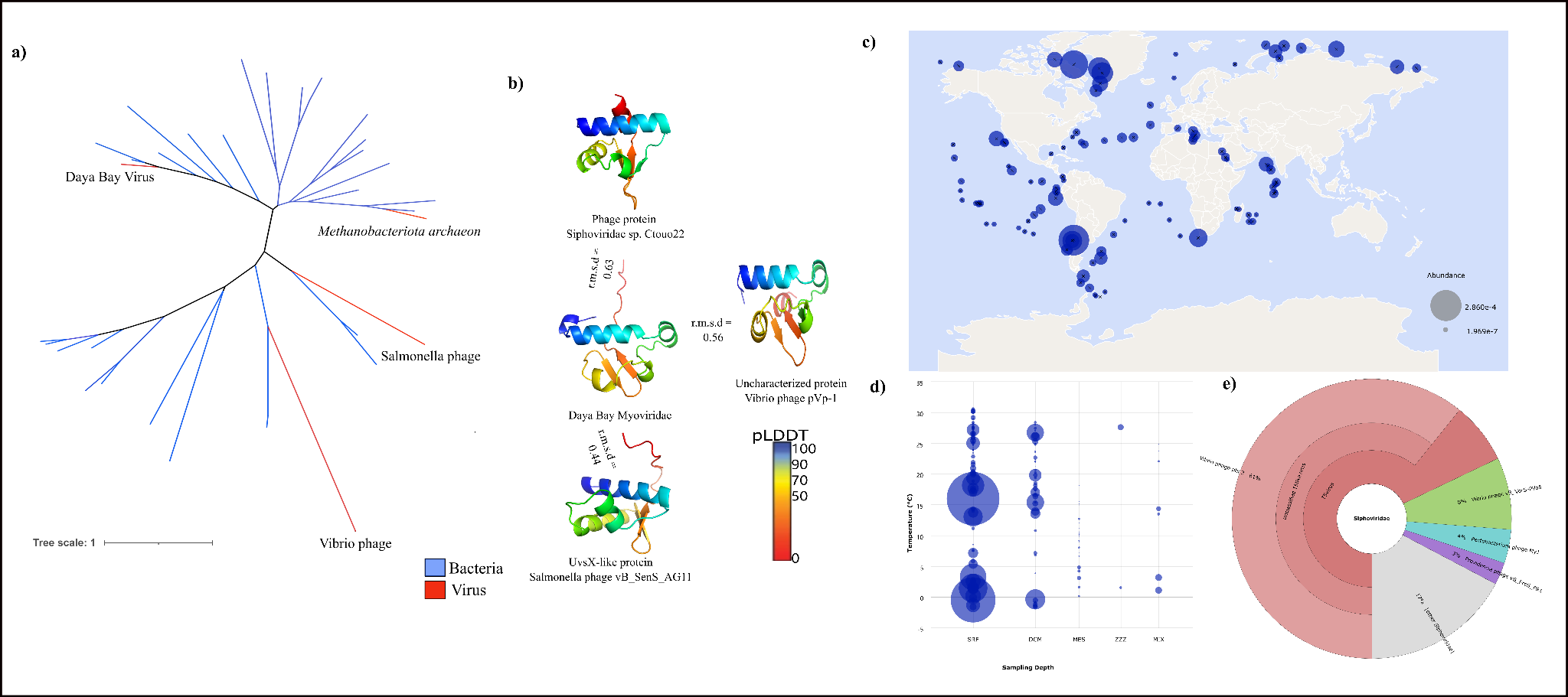

**Figure S9:** **Phylogeny, structure, and environmental distribution of a diel-cycling viral hypothetical gene**.

a) Maximum-likelihood phylogenetic tree of hypothetical protein sequences. The tree illustrates the evolutionary relationships of the viral hypothetical gene, showing significant diel patterns. Sequences include homologs from various viruses (red clade, including the highlighted "Daya Bay Myoviridae" and related phages) and bacterial hosts (blue clades). The scale bar represents the number of substitutions per site. (b) Predicted protein structure of Daya Bay virus and structural homologs. The AlphaFold3-predicted three-dimensional structure of the gene from the Daya Bay virus (middle) is shown. For structural comparison, *in silico* determined structures of homologous proteins from Siphoviridae sp*.* Ctouo22 (top), virbrio phage pVp-1 (right), and Salmonella phage vB_SenS_AG11 (bottom) are displayed. The alignment between these proteins is represented by root-mean-square distance (RMSD) values. The color gradient from blue to red represents the predicted Local Distance Difference Test (pLDDT) confidence score (blue: high confidence, red: low confidence). (c) Global abundance distribution of the gene. A world map depicting the global distribution of sequences homologous to the gene identified in this study. The size of the blue circles is proportional to the relative abundance of these similar sequences in publicly available metagenomic datasets from various marine environments. (d) Environmental distribution of the gene sequences across temperature and depth gradients. A bubble plot showing the abundance of gene (represented by bubble size) across different temperatures (°C) and sampling depth categories. Labels such as SAF (Surface Aphotic), ECH (Epipelagic Chlorophyll Max), MES (Mesopelagic), ZZZ (Zooplankton Rich Zone), and MIX (Mixed Layer) represent distinct marine water layers/regimes. This plot illustrates the prevalence of this gene across diverse environmental conditions. e) The corona plot shows the widespread distribution of this gene in similar Siphoviridae virus.

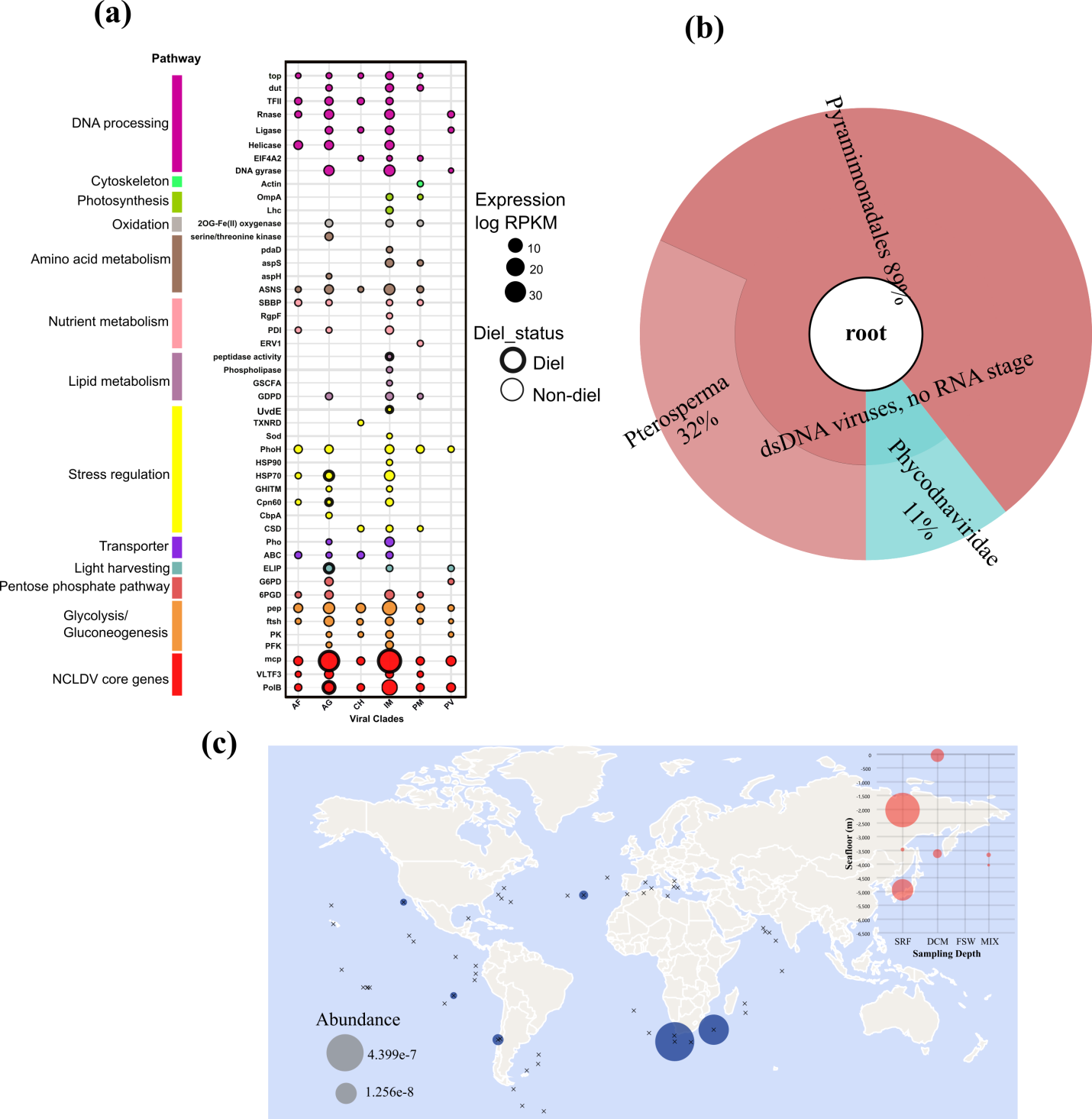

**Figure S10:** **Metabolic potential of the NCLDV and environmental distribution of a diel-cycling viral elip gene**. a) Metabolic genes of Daya Bay NCLDVs. The x axis shows different viral clades and the y axis denotes the functional annotation. The size of the bubble represent the total abundance of the gene in TPM (natural log transformed) and the colors show the functional category. Family assignment abbreviations: AF, Asfuvirales; IM, Imitervirales; AG, Algavirales; PM, Pimascovirales; and PV, Pandoravirales. Gene function abbreviations: top, Topoisomerase; tfs, Transcription factor; Rnase, RNase; EIF4A2, Eukaryotic Translation Initiation Factor 4A2; dut, dUTPase; ASNS, Asparagine synthase; aspH; Aspartate Beta-Hydroxylase; aspS, Aspartyl-tRNA synthetase; pdaD, Pyruvoyl-dependent arginine decarboxylase; PDIA, Protein disulfide-isomeraseA; GDPD, Glycerophosphodiester phoshodiesterase; UvdE. UV-damage endonuclease; TXNRD, Thioredoxin reductase; Sod, Superoxide dismutase; HSP, Heat shock protein; Cpn60, Chaperonin 60; CSD, cold shock domain; Pho; Phosphate regulator; ELIP, Early light-induced protein; G6PD, Glucose-6-phosphate dehydrogenase; 6PGD, 6-phosphogluconate dehydrogenase; Pep, peptidase; ftsh, ATP-dependent zinc metalloprotease; PK, protein kinase; mcp, major capsid protein; VLTF3, viral late gene transcription factor 3; PolB, polymerase beat; PFK, phosphofructokinase. Diel genes were represented with a thicker border. (c) Global abundance distribution of ELIP-like sequences. A world map depicting the global distribution of sequences homologous to the ELIP gene identified in this study. The size of the blue circles is proportional to the relative abundance of these ELIP-like sequences in publicly available metagenomic datasets from various marine environments. The bubble plot inside this graph shows the environmental distribution of ELIP-like sequences across temperature and depth gradients. This plot illustrates the prevalence of this gene across diverse environmental conditions. e) The corona plot shows the widespread distribution of this gene in similar NCLDVs.
